# Transposable element exaptation is the primary source of novelty in the primate gene regulatory landscape

**DOI:** 10.1101/083980

**Authors:** Marco Trizzino, YoSon Park, Marcia Holsbach-Beltrame, Katherine Aracena, Katelyn Mika, Minal Caliskan, George H. Perry, Vincent J. Lynch, Christopher D. Brown

## Abstract

Gene regulation plays a critical role in the evolution of phenotypic diversity. We investigated the evolution of liver promoters and enhancers in six primate species. We performed ChlP-seq for two histone modifications and RNA-seq to profile cis-regulatory element (CRE) activity and gene expression. The primate regulatory landscape is largely conserved across the lineage. Conserved CRE function is associated with sequence conservation, proximity to coding genes, cell type specificity of CRE function, and transcription factor binding. Newly evolved CREs are enriched in immune response and neurodevelopmental functions, while conserved CREs bind master regulators. Transposable elements (TEs) are the primary source of novelty in primate gene regulation. Newly evolved CREs are enriched in young TEs that affect gene expression. However, only 17% of conserved CREs overlap a TE, suggesting that target gene expression is under strong selection. Finally, we identified specific genomic features driving the functional recruitment of newly inserted TEs.

## Introduction

The evolution of cis-regulatory elements (CREs) plays an important role in phenotypic and behavioral evolution (King and Wilson, 1975; Rockman et al., 2005; Loisel et al., 2006; Pollard et al., 2006; Prabhakar et al., 2008; Warner et al., 2009; Babbitt et al., 2010; Cain et al., 2011; Marnetto et al., 2014; Zhou et al., 2014; Villar et al., 2015). Many aspects of CRE evolution in mammals have been characterized (Schmidt et al., 2010; Cain et al., 2011; Zhou et al., 2014; Prescott et al., 2015; Reilly et al., 2015; Villar et al., 2015; Emera et al., 2016), and suggest a role for transposable elements (TEs) in the evolution of gene regulation (McClintock, 1950, 1984; Britten and Davidson, 1969; Davidson and Britten, 1979; Jordan et al., 2003; Bejerano et al., 2006; Wang et al., 2007; Bourque et al., 2008; Sasaki et al., 2008; Markljung et al., 2009; Kunarso et al., 2010; Lynch et al., 2011, 2015; Chuong et al., 2013, 2016; Schmidt et al., 2012; Xie et al., 2013; del Rosario et al., 2014; Sundaram et al., 2014; Du et al., 2016; Rayan et al., 2016). However, validating the functional contribution of TEs in the mammalian gene regulation remains a challenge. Lynch et al. (2011, 2015) demonstrated that the recruitment of novel regulatory networks in the uterus was likely mediated by ancient mammalian TEs. However, Emera and colleagues (2016) suggested that neocortical enhancers do not exhibit strong evidence of transposon exaptation.

Many important questions remain unanswered: to what extent are poised and active regulatory elements functionally conserved? Are specific genomic features predictive of CRE conservation? To what extent have TEs driven the evolution of gene regulation? And finally, what determines which TE insertions are recruited as functional CREs? Establishing answers to these questions is critical for understanding how the evolution of regulatory elements contributes to the conservation and divergence of gene expression and complex traits.

With a goal of answering these questions, we collected liver samples from six primate species. Core liver functions are largely conserved across primate species. However, different environmental exposures, diets, and lifestyles likely directed the adaptation of liver functions, and associated regulatory evolution, making this tissue an optimal model in which explore the conservation and divergence of the gene regulation.

To characterize primate liver cis-regulatory evolution, we performed chromatin immunoprecipitation followed by sequencing (ChlP-seq) for Histone H3 Lysine 27 acetylation (H3K27ac), which marks active enhancers and promoters, and Histone H3 Lysine 4 mono-methylation (H3K4me1), which marks poised regulatory elements on liver tissues from six primate species, including at least one species from each major primate clade to maximize evolutionary diversity within primates (Perelman et al., 2012). We generated whole transcriptome sequencing (RNA-seq) data from the same specimens to quantify gene expression variation across species. We estimated the degree of evolutionary conservation of regulatory activity and gene expression levels across the entire lineage, and characterized the genomic features associated with evolutionary conservation of gene regulation, to understand why some enhancers and promoters are conserved across species whereas others are subject to rapid turnover.

The activity of the majority of human CREs is conserved across the entire primate lineage, and the differences in gene expression and regulation reflect the phylogenetic distance between species. Conservation of cis-regulatory activity is associated with nucleotide sequence conservation, gene function, gene distance, cell-specificity of CRE function, and transcription factor binding site (TFBS) density. Strikingly, human- and ape-specific enhancers and promoters are significantly enriched for evolutionarily young TEs. In particular, the majority of human- and ape-specific CREs are derived from SINE-VNTR-*Alus* (SVAs) and Long-Terminal-Repeats (LTRs), respectively. On the other hand, only a minor fraction of evolutionarily conserved CREs are derived from TEs, indicating that purifying selection on the associated genes likely preserve these CREs from being disrupted by TE insertions, thus conserving the expression of the associated genes. In addition, we characterized SVAs that evolved into regulatory elements, and estimated potential impacts of these SVAs on gene regulation across the lineage. We validated the regulatory activity of several TE families, and conclude with a new model describing specific genomic features that strengthen the potential adaptation of TEs into functional regulatory elements (exaptation; Brosius and Gould; 1992; de Souza et al., 2013).

## Results

### Data generation, quality assessment, and validation

We generated a total of 757 million RNA-seq reads and 1.70 billion ChIP-seq reads (H3K27ac, H3K4me1, and input) from *post mortem* livers of three or four individuals per species of mouse lemur (*Microcebus murinus*), bushbaby (*Otolemur garnettii*), marmoset (*Callithrix jacchus*), rhesus macaque (*Macaca mulatta*), chimpanzee (*Pan troglodytes*), and human (*Homo sapiens*) (Fig. 1; Table S1). The six species were selected to include at least one species from each of the major primate clades, thus maximizing phylogenetic diversity within primates. The samples were all from young adults and, with the exception of the bushbaby, included both males and females. In total, 18 RNA-seq and 14 ChIP-seq samples remained post-QC and were used for analyses (Table S1). On average, we sequenced 42.1 million RNA-seq reads and 40.6 million ChIP-seq reads per sample (Table S1). We applied stringent quality control (QC) measures to assess library construction, sequencing, and peak-calling methods. Read mappability after the filtering steps was consistent across species and assays (77% in ChIP-seq, 78% in RNA-seq; Table S1), suggesting that differences in genome assembly quality do not introduce large biases.

**Figure 1 -.**
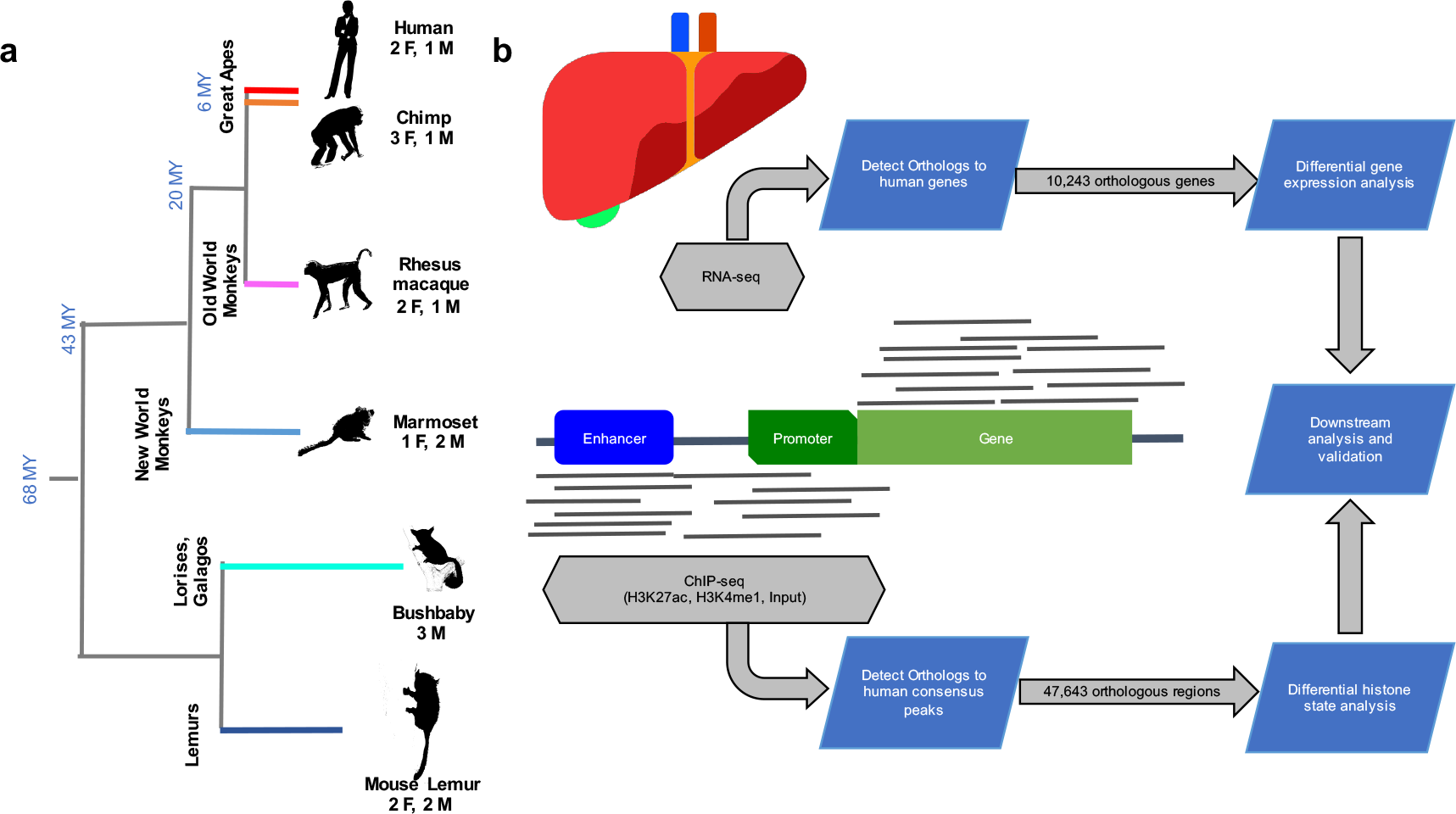
Experimental design and analytical pipeline. (A) Sampling included three to four specimens from six species representing all of the major primate clades. (B) ChlP-seq and RNA-seq profiles were produced from the liver samples. Differential histone modification and gene expression analyses were performed on the orthologous CREs and genes in each species, respectively. Outputs shown in the diagram were used for downstream analyses and validations.

ChIP-seq data were used to map the distribution of active (H3K27ac) and poised (H3K4me1) CREs in the six primate genomes (Fig. 1). We mapped ChIP-seq reads to their respective reference genomes with BWA-mem (Li, 2013) and identified regions of significant H3K27ac and H3K4me1 enrichment in the human liver, treating all human individuals as replicates in the peak calling procedure with MACS2 (Zhang et al., 2008). 85.6% of the regions marked by H3K4me1 overlapped regions marked by H3K27ac. A total of 84,253 human peaks remained after merging overlapping peaks from the two histone markers. Next we identified regions orthologous to the human consensus peaks from the genomes of non-human primates using the Ensembl multiple sequence alignment (MSA) database. We catalogued 47,673 total human CREs with orthologs in all six species: 40,527 enhancers (distance from transcription start site (TSS) > 1 kb) and 7,146 promoters (distance from TSS ≤ 1 kb).

Several lines of evidence indicate that the regions of histone modification we have identified represent active CREs. First, 99.0% and 73.3% of ENCODE HepG2 H3K27ac and H3K4me1 regions (ENCODE Project Consortium, 2012), respectively, overlapped with one or more human peaks. Similarly, 63.2% of the 43,011 permissive enhancers that were predicted by the FANT0M5 Consortium based on enhancer RNA expression (Andersson et al., 2014) overlapped with one or more of human peaks identified in our study. Further, the promoters of 98.1% of genes expressed in the liver overlapped a region of histone modification (Fig. S1). Finally we compared our human H3K27ac data to a recent study focused on liver CREs in mammals (Villar et al., 2015) and demonstrated that peaks bearing signatures of robust and broad regulatory activities are largely reproducible across studies, despite variation attributable to different study designs (Fig. S2).

ChlP-seq experiments can identify regions of the genome bound by histones and other proteins that characterize regulatory elements, however, this does not guarantee these regions are functional regulatory elements (Pickrell et al., 2011; Cusanovic et al., 2014; Jain et al., 2015). Therefore, we used a novel parallelized reporter assay (Melnikov et al., 2012; Patwardhan et al., 2012; Sharon et al., 2012; see methods) to validate the regulatory function of predicted human liver CREs. Specifically, we tested the regulatory activity of 1 kb fragments from 122 putative regulatory elements in HepG2 cells, including both evolutionarily conserved and recently evolved CREs (see below). Among 122 tested elements, 79 drove significant reporter gene expression levels (42/53 enhancers [79.2%] and 36/69 promoters [52.2%]; Table S6), suggesting that the majority of the CREs predicted in our study based on the enrichment of active histone modification states are likely functional regulatory elements in the human liver.

### The majority of human CREs are functionally conserved across primates

After extracting ChIP-seq read counts for the six species from 47,673 orthologous regions, we assessed evidence of differential histone modification between species with DESeq2 (Love et al., 2014), using the ChIP-seq input data as a covariate. We compared ChIP-seq read counts in the 47,673 regions by means of all possible human-centric species × species and group × group pairwise comparisons (see methods). This approach provides a quantitative assessment of histone modification profiles across species, while avoiding issues arising from many potential experimental variables that may confound peak calling (Waszak et al., 2015). A specific analysis of human and marmoset, the latter being the species with the smallest number of peaks called in this study (Supplemental File S1), strongly supported the validity and robustness of our approach (Fig. S3).

The majority of the 47,673 human CREs (63.8%) did not show significant differential histone modifications in any of the tested pairwise comparisons (FDR < 10%; Fig. 2). This suggests that these regulatory regions are consistently active across the primate lineage and thus represent evolutionarily conserved primate CREs. Although the absence of differential histone modifications in a pairwise comparison between two species is not a direct proof of CRE conservation, we demonstrated that the selected FDR threshold does not affect downstream conclusions (Table S7). As an additional control, we performed a chimpanzee-centric analysis for the regions orthologous to chimpanzee consensus peaks (hereafter, chimpanzee CREs), and demonstrated that 62.5% of these regions were not differentially histone modified in any of the pairwise comparisons. This observation is consistent with 63.8% conserved CREs identified in the human-centric analysis, indicating that the differential histone modification analysis is robust and species-specific bias is unlikely.

**Figure 2 -.**
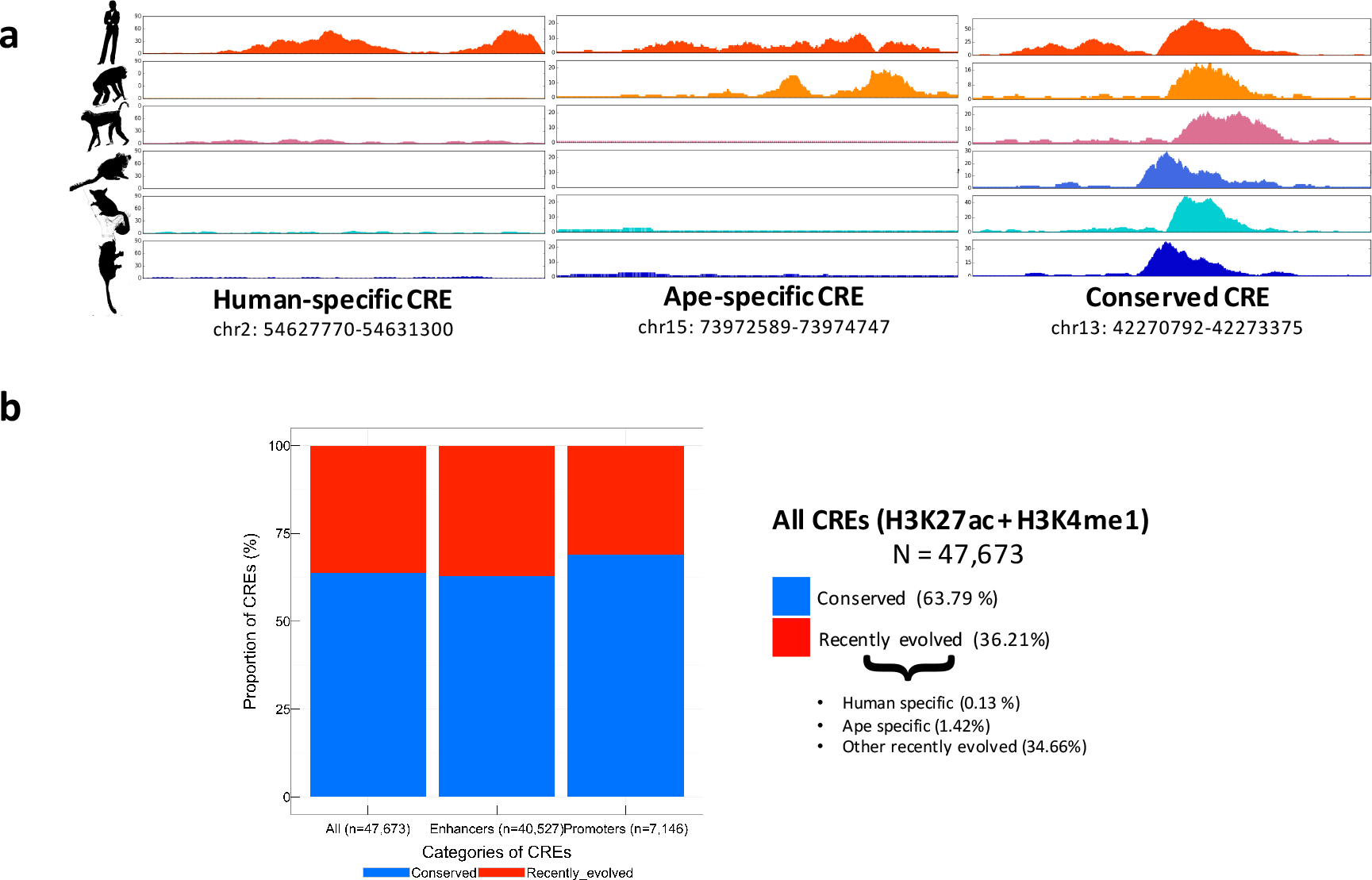
Primates CREs are evolutionarily conserved. (A) Plots show examples of human-specific, ape-specific and conserved CREs. (B) Fractions of conserved and recently evolved primate CREs, with breakdown of enhancers and promoters.

Promoters are significantly more conserved than enhancers (68.9% and 62.8%, respectively; Fisher’s exact test *p* < 2.2×10^−16^; Fig. 2), as observed previously (Villar et al., 2015). On the other hand, 36.2% of orthologous CREs exhibited differential histone modification state across species (Fig. 2). We detected 57 human-specific CREs (0.13%; Fig. 3) and 544 ape-specific CREs (1.42%; Fig. 3). Together, our differential histone state analysis results are broadly supported by several studies that have consistently suggested a high degree of regulatory element conservation between closely related species in metazoans (Cotney et al., 2013; Boyyie et al., 2014; Prescott et al., 2015; Reilly et al., 2015; Emera et al., 2016).

**Figure 3 -.**
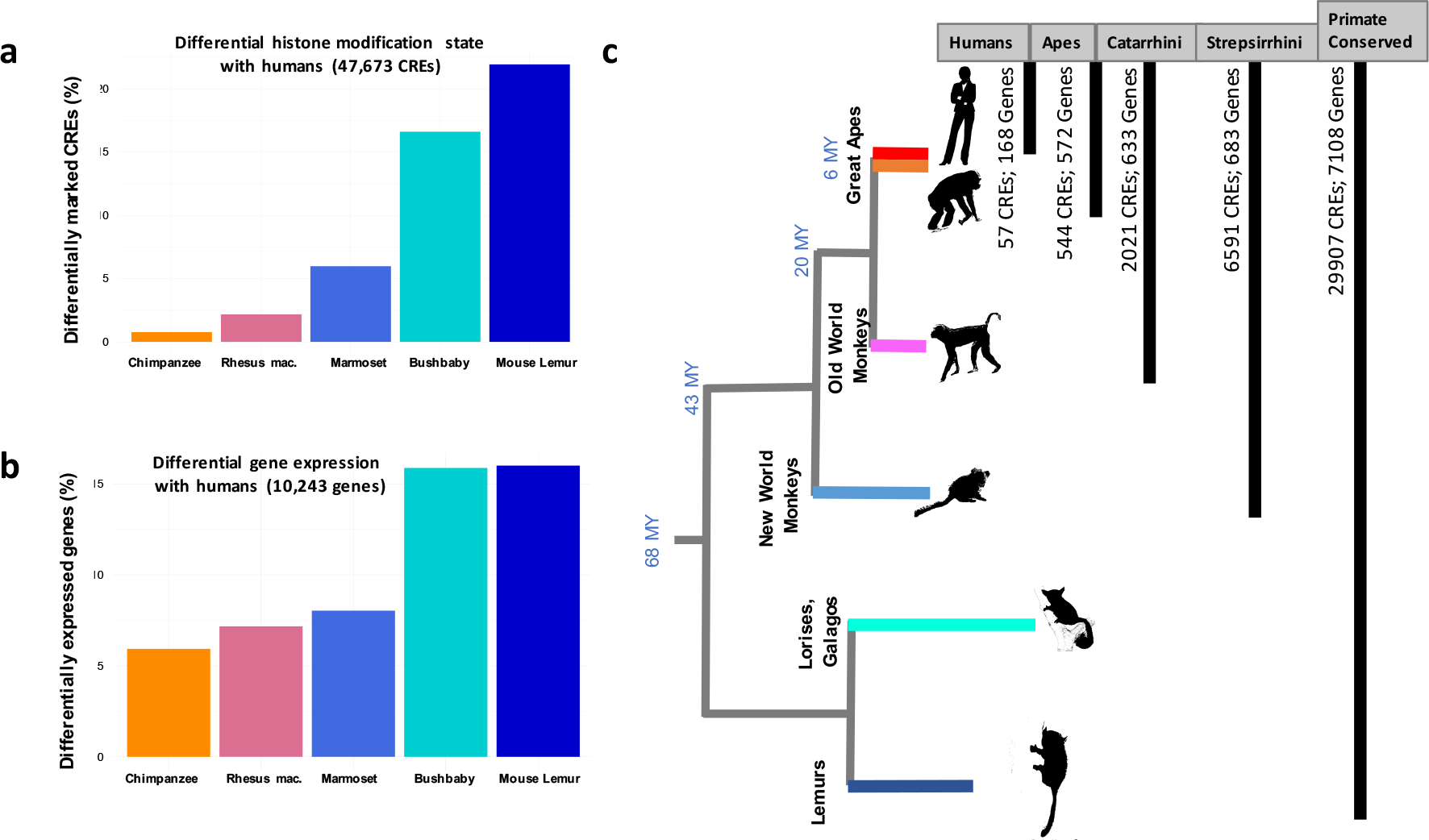
Differential histone mark and gene expression across species. (A) Human-centric pairwise comparisons for differential histone modification states on 47,673 orthologous CREs. (B) Human-centric pairwise comparisons for differential gene expression of 10,243 orthologous CREs. (C) Number of lineage-specific CREs (i.e. CREs significantly more active in each lineage compared to other primates) and genes (i.e. genes upregulated in each lineage compared with other primates) in the primate phylogeny.

Next, we assessed the extent to which primate-conserved CREs identified in this study are also evolutionarily conserved across a broader range of mammals. In particular, we compared our conserved H3K27ac CREs with the H3K27ac profile of the opossum, the species with the earliest divergence from humans (>180 million years) in the Villar et al. (2015) dataset. 2,854 primate-conserved promoters and 9,456 primate-conserved enhancers have orthologous regions in the opossum genome. Among these, 71.3% of the promoters and 19.1% of the enhancers had significant H3K27ac enrichment in both species, supporting that most of the primate conserved promoters show conserved activity in all of the mammalian clade, whereas only a fraction of the primate conserved enhancers is also conserved across mammals. Further, the two studies come to consistent estimates of the fraction of differentially active CREs per million years: 0.06–0.12% in primates and 0.07% in mammals.

### The conservation of the nucleotide sequence is associated with conservation of regulatory activity

Previous studies have suggested that the sequence conservation is associated with conservation of regulatory activity, especially in absence of comparative functional assays (Brown et al., 2007; Cooper and Brown, 2008; Pollard et al., 2010; Gittelman et al., 2015; Holloway et al., 2015; Villar et al., 2015; Yang et al., 2015; Dong et al., 2016; Lewis at al., 2016). For each human-centric species × species comparison, we estimated: i) the fraction of differentially modified CREs; ii) the fraction of differentially expressed genes from a set of 10,243 genes with six way orthologs (Table S3); and iii) the per-nucleotide pairwise sequence divergence for each species with respect to humans for each of the 47,673 unique orthologous CREs

Differential histone state fractions ranged from 0.77% in the human × chimpanzee, to 21.9% in the human × mouse lemur comparisons (Fig. 3a). Similarly, differential gene expression ranged from 5.93% in the human × chimpanzee to 16.0% in the human × mouse lemur comparisons (Fig. 3b). Both differential histone modification and differential gene expression fractions reflected phylogenetic distance between humans and other tested species. Differentially expressed genes were significantly more likely to be associated with a differentially modified CRE than expected by chance (9.90%; Fisher’s exact test *p* < 2.2×10^−16^).

Sequence conservation was significantly correlated with regulatory activity (human × chimpanzee, logistic regression *p* = 5.7×10^−16^; human × rhesus macaque, *p* = 6.7×10^−8^; human × marmoset, *p* = 4.0×10^−9^; human × bushbaby, *p* < 2.2×10^−16^; human × mouse lemur, *p* < 2.2×10^−16^; Fig. 3c). 24,691 CREs overlapped 94,578 placental mammal phastCons elements (i.e. regions of the genomes with consistent nucleotide sequence conservation across species; Siepel et al., 2005). The fraction of evolutionary conserved CREs overlapping these conserved elements was higher than expected by chance (Fisher’s exact test *p* < 2.2×10^−16^). Together, these data demonstrate that CREs with conserved nucleotide sequence are significantly more likely to have conserved regulatory activity and are associated with conserved gene expression.

### Genomic features associated with CRE conservation and rapid evolution

To understand the mechanisms responsible for CRE conservation and turnover, we identified genomic features associated with conserved regulatory activity. CREs associated with protein coding genes were significantly more conserved than CREs associated with either pseudogenes (Fisher’s exact test *p* = 3.3×10^−3^) or lincRNAs (Fisher’s exact test *p* = 5.2×10^−7^; Fig. 4a). For closely related species, regulatory activity was conserved, regardless of the distance to the nearest TSS (human × chimpanzee, logistic regression *p* = 0.261; human × rhesus macaque, *p* = 0.336; Fig. 4b). However, for more distantly related species pairs, the evolutionary conservation of the CRE activity was significantly lower in regions more distant from TSSs (human × marmoset, logistic regression *p* = 6.0×10^−15^; human × bushbaby, *p* = 2.1×10^−5^, human × mouse lemur, *p* = 2.8×10^−5^; Fig. 4b). Intronic enhancers were significantly more conserved than intergenic enhancers (63.0% and 55.2% respectively; Fisher’s exact test *p* < 2.2×10^−16^). These data demonstrate increased selective pressure to maintain regulatory activity in the vicinity of protein coding genes.

**Figure 4 -.**
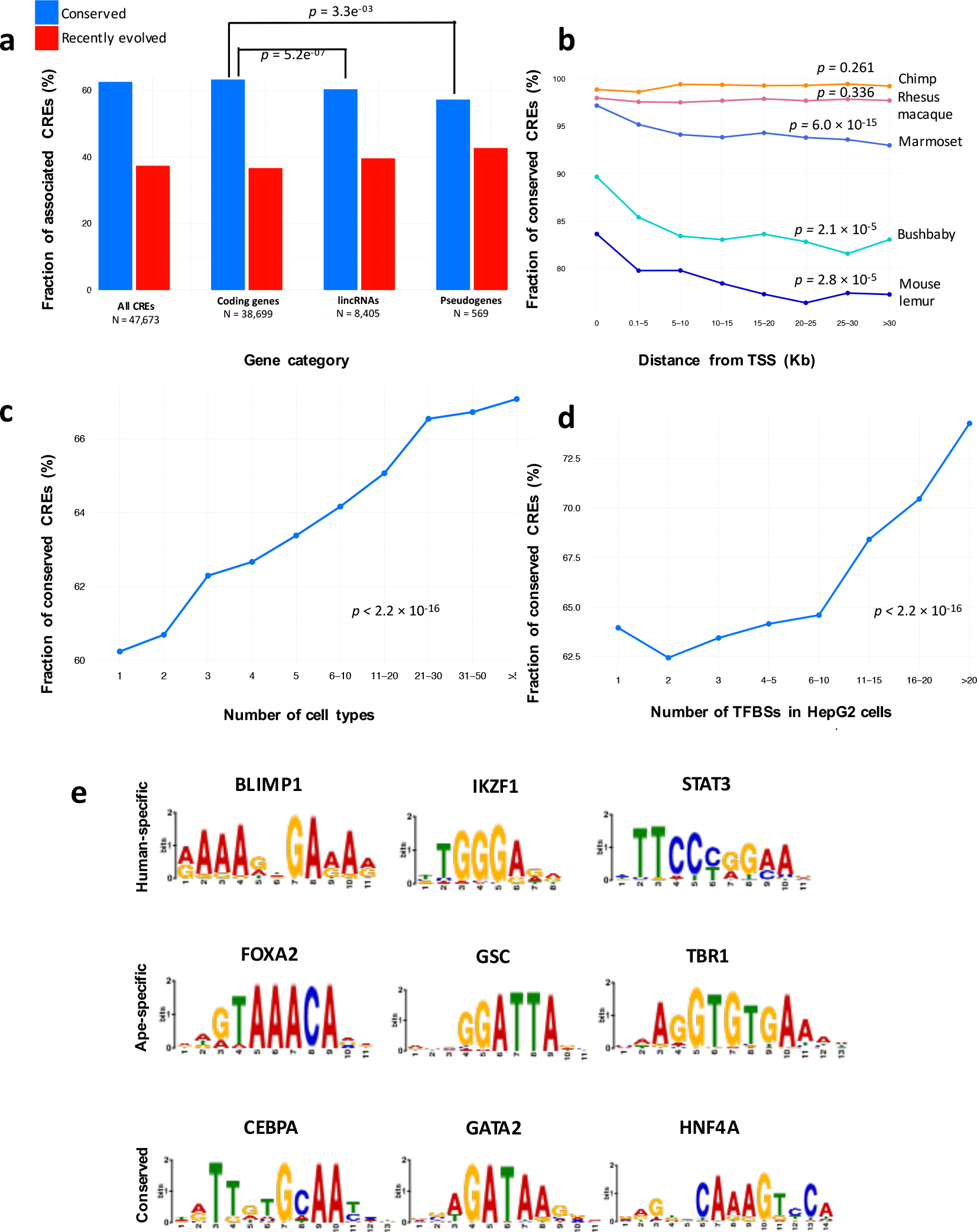
Genomic features associated to CRE conservation. (A) Fractions of conserved and recently evolved CREs associated to protein coding genes, lincRNAs and pseudogenes. (B) Distribution of CRE conservation along the genome. (C) Correlation between CRE conservation and cell-type specificity based on ENCODE data. (D) Correlation between CRE conservation and number of ENCODE HepG2 TFBSs present of each CRE. (E) Examples of i) enriched motifs associated to immune response (BLIMP1, IZKF1, STAT3) in human-specific CREs; ii) enriched motifs associated to language and neural development (FOXA2, GSC, TBR1) in ape-specific CREs; iii) enriched motifs associated to master regulators (CEPBA, GATA2, HNF4A) in conserved CREs.

Multiple genomic features indicative of broad regulatory element activity were significantly associated with the conservation of regulatory activity across the primate phylogeny. Promoters and enhancers overlapping regions of chromatin accessibility in many cell types (ENCODE Project Consortium, 2012) were significantly more functionally conserved than those that are functional in only a small number of tissues (logistic regression *p* < 2.2×10^−16^; Fig. 4c). Similarly, CREs that overlapped many TFBS, as identified by ENCODE ChIP-seq in HepG2 cells, were significantly more evolutionarily conserved than those with fewer binding sites (logistic regression *p* < 2.2×10^−16^; Fig. 4d). Further, liver enhancers that generate consistent enhancer RNA transcription were significantly more evolutionarily conserved than the untranscribed enhancers (Fig. S3; Fisher’s exact test *p* = 0.04613), validating previous findings (Andersson et al., 2014).

Finally, we used GOrilla (Eden et al., 2007; Eden et al., 2009) to identify biological processes Gene Ontology terms that are enriched in genes found within 10-kb of evolutionarily conserved CREs, using as background all of the genes found within 10 kb from any of the 47,673 orthologous CREs. We found an enrichment for housekeeping functions involved in the regulation of cellular, transcriptional and developmental processes (Table S4). These findings support previous observations that conserved CREs are proximal to housekeeping genes (FANTOM5 Consortium, 2014; Villar et al., 2015).

### Specific transcription factor motifs are associated with regulatory conservation and turnover

We used the MEME Suite (Bailey et al., 2009) to identify sequence motifs enriched in human-specific, ape-specific, and evolutionarily conserved liver CREs. Human-specific CREs were enriched with motifs for TFs associated with immune response and hematopoietic maintenance (Fig. 4e; Supplemental File S2), such as PRDM1 (BLIMP1) which is induced upon viral infection and represses beta-interferon (β-IFN) gene expression. The rapid evolution of immune response genes and TFs is supported by many studies in vertebrates and in *Drosophila melanogaster*, demonstrating that while the central machinery of immune responses is strongly conserved, several components of the extended molecular networks can evolve rapidly or diversify as a consequence of evolutionary competition between hosts and pathogens (Jansa et al., 2003; Vallender 2004; Sackton et al., 2007; Obbard et al., 2009; Schadt et al., 2009; Grueber et al., 2014; Lazzaro and Schneider 2014; Salazar-Jaramillo et al., 2014; Zak et al., 2014; Sironi et al., 2015; Wertheim, 2015; Chuong et al., 2016). Similarly, recently evolved promoters and enhancers in primates are enriched in functions associated to neuronal proliferation, migration and cortical map organization (Boyd et al., 2015; Reilly et al., 2015; Emera et al., 2016). In contrast, ape-specific CREs were instead enriched with motifs representing binding sites for TFs involved in liver function but, remarkably, also in brain and neural system proliferation and development (Fig. 4e; Supplemental File S2).

Evolutionarily conserved CREs were enriched in TFBSs for master regulators and homeobox genes that establish cell-type identity in liver cells (Fig. 4e; Supplemental File S2). Among these master regulators, HNF4A is essential for the differentiation of human hepatic progenitor cells by establishing the expression of the network of transcription factors that controls the onset of hepatocyte cell fate (DeLaForest et al., 2011). Likewise, CEBPA is required for the liver cell specification and gene function, and the associated TFBSs are highly conserved across mammals (Ballester et al., 2014). Both CEBPA and HNF4A have conserved cis-regulatory activity and significantly higher numbers of shared TF binding events than expected by chance alone across distant vertebrates (Schmidt et al., 2010). These results demonstrate that evolution shapes the regulatory landscape by preserving the regulatory activity in essential metabolic and developmental pathways, while permitting incessant renovation of specific networks that are under strong selective pressures.

### Exaptation of TEs into functional CREs is pervasive in the primate genomes

Previous studies have found that TEs can contribute to the origin of CREs. To quantify the contribution of TEs in the liver gene expression regulation in primates, we annotated each liver CRE based on overlap with RepeatMasker elements (Smit, Hubley & Green, 2013-2015; http://www.repeatmasker.org). We found that 28.7% of human liver CREs overlapped an annotated TE, most of which were SINEs (59.8% of the total) and LINEs (21.2%), although LTRs (9.3%) and DNA transposons (8.4%) were also abundant. 27 TE families were significantly enriched within the set of 47,673 orthologous CREs (FDR < 1%), nearly all of which were SINE-VNTR-Alus (SVAs), LTRs, and *Alus* (Figure 5), suggesting these TEs contributed to the regulatory landscape in the primate liver (Jordan et al., 2003; Schmidt et al., 2012; Sundaram et al., 2014; Lynch et al., 2015).

**Figure 5 -.**
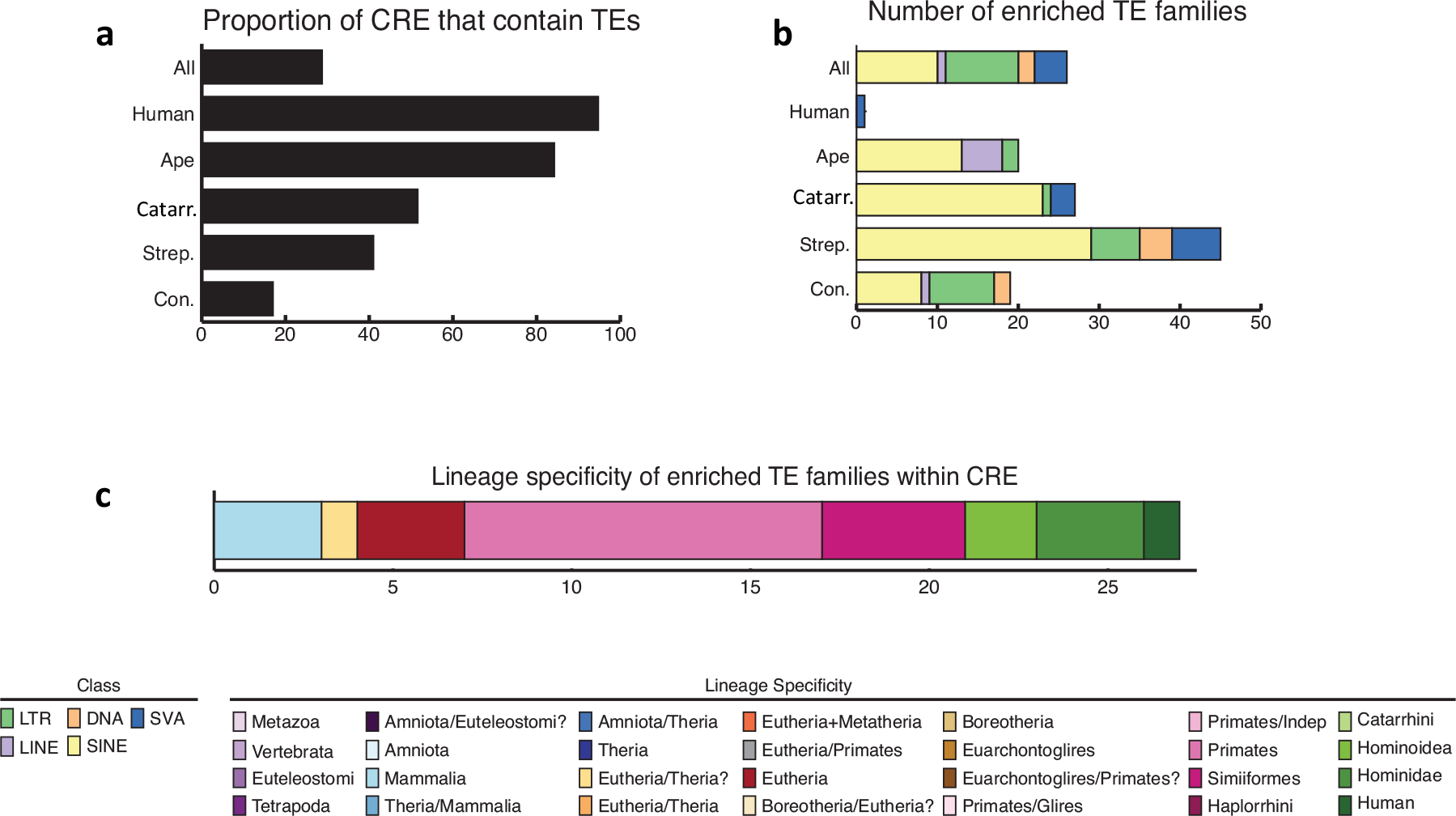
Newly evolved CREs areenriched in TEs. (A) Proportion of CREs that overlap TEs in the different primate lineages. (B) Number of enriched TE families within CREs in the different primate lineages. (C) Most enriched TE families in primates.

The majority (75.0%) of the enriched TE families were relatively young, and specific to humans (SVA-F), Hominidae (SVA-B, SVA-C, and SVA-D), Hominoidea (the LTR12 subfamily), Simiiformes (LTRs), or primates (Alu elements), whereas the remaining 25.0% were Eutherian-specific or older (Fig. 5, Fig. S7, and Table S5). We therefore investigated whether these recent TE insertions altered the expression patterns of nearby genes in primates. Specifically, we focused on the enriched TE families younger than the Strepsirrhini/Haplorrhini divergence, thus not present in mouse lemur and bushbaby. We found that 22.6% of the CREs overlapping SVAs, and 12.5% of the CREs overlapping LTRs were differentially expressed between the Strepsirrhini (human, chimpanzee, rhesus macaque and marmoset) and the Haplorrhini (mouse lemur and bushbaby). Both of these fractions were significantly higher than expected by chance (binomial test *p* < 2.2×10^−16^). Together, these findings indicate that TEs have played a key role in shaping primate gene regulation, introducing novel gene expression patterns as a consequence of their recruitment as functional CREs.

### The vast majority of newly evolved CREs are derived from TE insertions

84.2% of ape-specific CREs and 94.7% of human-specific CREs overlap at least one TE (Fig. 5; Fig. S7). In contrast, only 17.0% of evolutionarily conserved CREs contain an annotated TE (Fisher’s exact test *p* < 2.2×10^−16^). However, we hypothesize that this may be an underestimate due to the inability to recognize ancient TE insertions accumulating mutations over time. LTRs and SVAs were the most frequently exapted TEs in newly evolved CREs (LTR = 40.1% of the exapted TEs in ape-specific CREs; SVA = 75.3% of the exapted TEs in the human-specific CREs; Fig. 5), despite being among the least common classes of repeats in the human genome (15.9% and 0.69% of the total TEs respectively; Fisher’s exact test *p* < 2.2x10^−16^ for both of the TE categories).

The contribution of LTRs in primate gene regulation has been characterized in previous studies (Wang et al., 2007; Cohen et al., 2009; Sundaram et al., 2014; Janousek et al., 2016). In our data, a remarkable example of an ape-specific CRE derived from LTR insertion is an enhancer at the gene *GRIN3A.* This gene is involved in physiological and pathological processes in the central nervous system and has been associated with several complex human diseases, including schizophrenia (Takata et al., 2013). Our differential histone modification analysis identified an ape-specific ChlP-seq peak overlapping a 1-Kb long ape-specific insertion (present also in orangutan and gorilla, but not in other primates; GRCh38 chr9:101,723,127-101,724,197). This insertion, located 13 Kb from the TSS of *GRIN3A*, is entirely derived from an LTR-12C. The insertion drove strong enhancer activity upon transfection into HepG2 cells (Wilcoxon’s rank sum test *p* = 0.00017; Fig. S5), suggesting that the transposable element was recruited as functional enhancer in the *GRIN3A* locus.

SVAs are instead a hominid-specific family of composite retrotransposons that are active in humans (Hancks and Kazazian, 2010), with more than 3,500 annotated copies. Given that nearly all human-specific liver CREs were derived from SVA insertions in our analysis, we further investigated the genomic features of SVA insertions that lead to exaptation (Table S5). 49.5% of human SVAs overlapped regions of significant histone modification, and 97.8% of those were enhancers. These exapted SVAs are significantly more likely to be associated with protein coding genes than the non-exapted SVAs (Fisher’s exact test *p* = 0.017) and are significantly closer to the TSS of the associated gene (mean of 52.9 kb versus 64.1 kb; Wilcoxon rank-sum test *p* < 2.2×10^−16^). Exapted and non-exapted SVAs lie within open chromatin regions in approximately the same number of cell types (3.44 and 3.94 respectively; logistic regression *p* = 0.827) and host on average a comparable density of TFBSs (3.28 and 1.95; logistic regression *p* = 0.679). However, exapted SVAs have a significantly higher number of TFBSs in the neighboring regions (8.12 versus 5.86 in +/− 10kb; logistic regression *p* = 1.91×10^−5^). Taken together, these data suggest a model where an SVA has a higher probability of becoming a CRE if it inserts in TFBS-dense regions near protein coding gene promoters. (Fig. 6).

**Figure 6 -.**
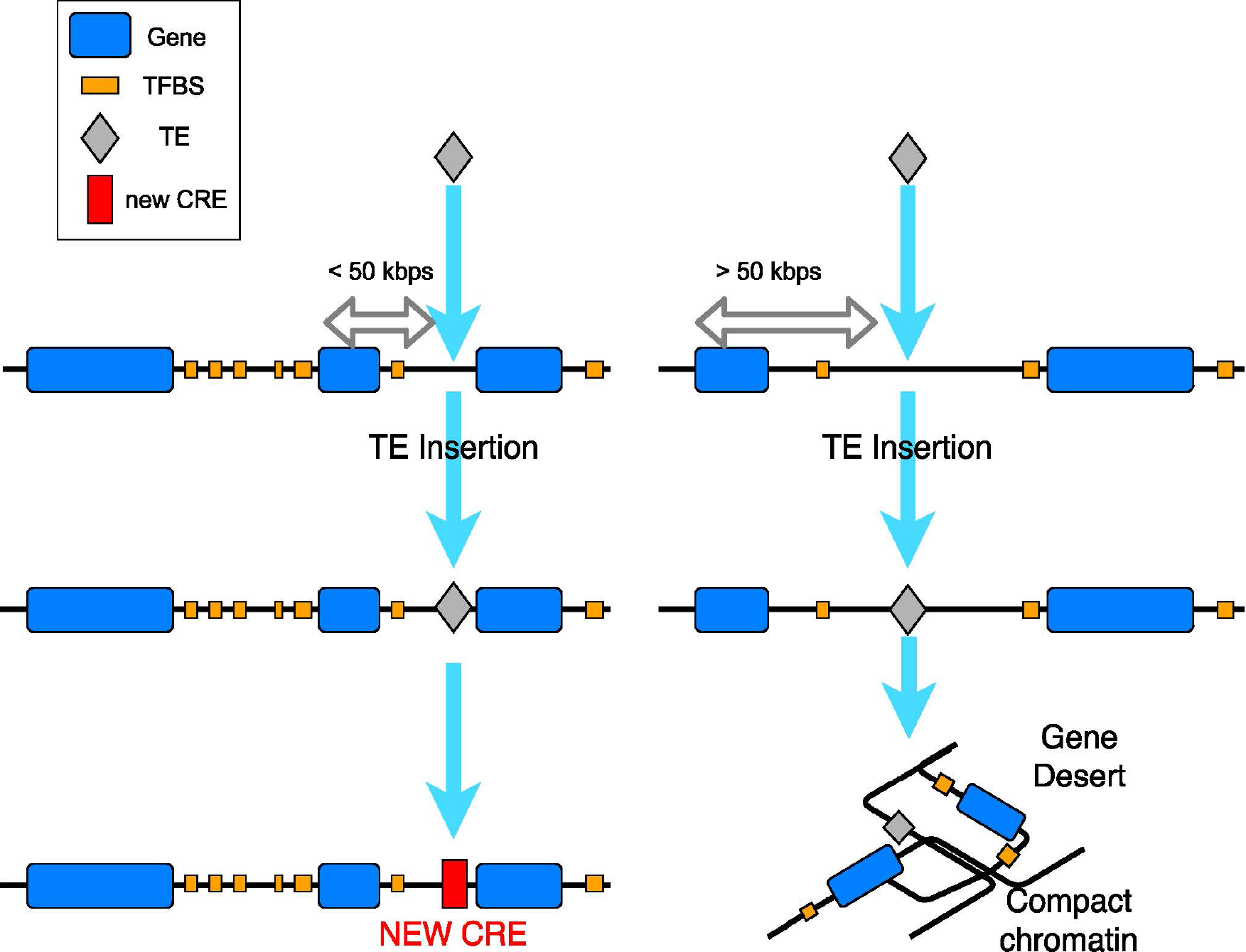
Model for exaptation of SVAs into human functional CREs. A given SVA has a higher probability of being recruited as a functional cis-regulatory element if it is found within 50 kb from a protein coding gene, in a genomic region already enriched in TF binding motifs in the surrounding 10 kb up- and downstream.

Among the exapted SVAs, our data predicted an intronic CRE for the gene *JARID2.* This gene is an accessory component of Polycomb Repressive Complex-2 (PRC2), recruits PRC2 to chromatin, and is involved in liver, brain, neural tube development, and embryonic stem cell differentiation (Kaneko et al., 2014). Our differential histone state analysis revealed a human-specific ChIP-seq peak overlapping a human-specific 1.9 kb-long insertion, entirely derived from an SVA-F retrotransposon. JARID2 is significantly downregulated in humans compared to all the other primates (Benjamini–Hochberg *p* = 0.019). Exapted SVAs-Fs exhibit significant enrichment for binding sites of known transcriptional repressors such as PAX-5, FEV, and SREBF1 (Maurer et al., 2003; Fazio et al., 2008; Lecomte et al., 2010). Indeed, the JARID2 SVA-F insertion leads to significantly decreased expression in HepG2 reporter assays (Wilcoxon’s rank sum test *p* = 0.00275; Fig. S6), supporting the role of this SVA-F as a transcriptional repressor.

### Broad regulatory activity of TE insertions in the primate liver

Our findings strongly suggest that the majority of novel CREs in primates is derived from TE insertions. To validate the predicted regulatory activity of recent TE insertions, we tested the cis-regulatory activity of 69 TE subfamilies, covering all of the main classes and families of TEs (Table S6). TEs from these families overlap 3,897 of our predicted CREs. We synthesized the mammalian consensus sequence (see experimental procedures) for 69 different TE families, cloned them into a luciferase reporter vector with a minimal promoter, and transfected them into HepG2 cells to perform dual luciferase reporter assays. We found that luciferase expression for 66 of the 69 (95.6%) tested TE families was significantly different from the negative control (Fig. 7a; Wilcoxon’s rank sum test p-values in Table S6). Strikingly, only 17 (25.7%) of these, mostly LTRs and DNA transposons, produced activity significantly higher than the negative control (Fig. 7a), whereas the remaining 49 (74.3%), mostly LINEs, repressed transcription. Consistent with results from the JARID2 locus presented above, SVA-Fs were confirmed to function as transcriptional repressors.

**Figure 7 -.**
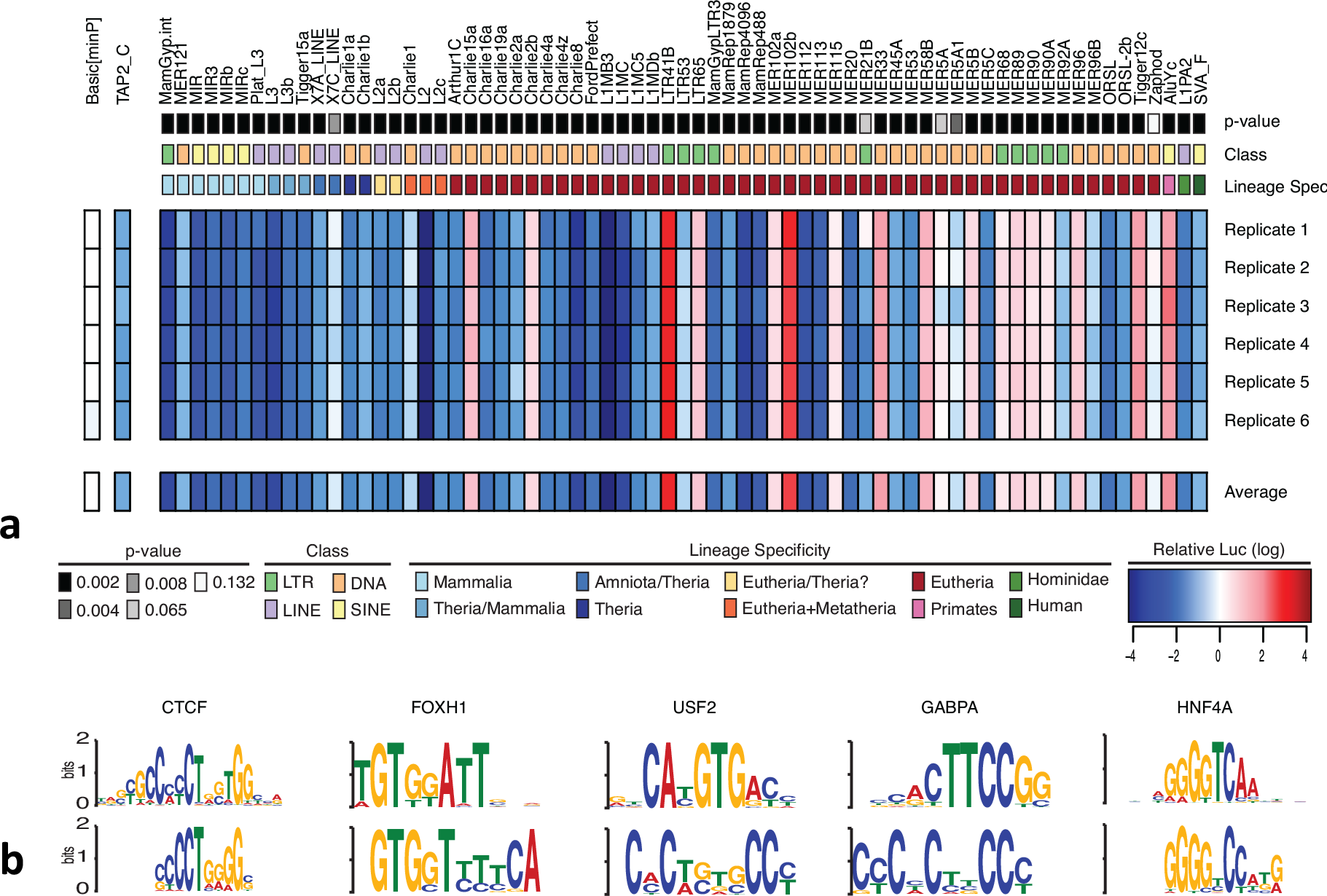
Regulatory ability of TE families found in poised and active regulatory elements in HepG2 cells. (A) The p-value, class, and lineage specificity for the 69 TE families tested in HepG2 cells for regulatory ability, the empty vector control (Basic[minP]), and the positive control (TAP2_C) are all shown above the 6 luciferase assay replicates conducted and the average regulatory ability found across the replicates. Red indicates luciferase expression higher than the empty vector control; blue indicates luciferase expression less than Basic[minP]. (B) CTCF, FOX, USF2, GABP, and HNF4a binding motifs were found to be enriched in the sequences of the 66 TE families that drive expression significantly different from background. The top row shows the enriched sequence found while the bottom shows the Jaspar binding motifs recognized for each transcription factor.

These findings support that LTRs, among the most enriched TEs in our peak set, and the most common exapted TEs in apes, are likely co-opted as active enhancer elements. The putative repressor activity of LINEs is consistent with their underrepresentation in the human ChIP-seq peak set. However, we hypothesize that, at least for some of the TE families, such observed repressing activity levels could be the byproduct of secondary biological mechanisms leading the cell to recognize these elements as newly inserted TEs, and therefore silencing them via epigenetic mechanisms. However, further tests will be needed to support this hypothesis.

The consensus sequences for the 66 TEs that drove reporter expression significantly different from background were analyzed with MEME to identify enriched motifs. Motifs for known master regulators of liver cell identity, including FOX, USF2, GABP, and HNF4A (Wallerman et al., 2009), were significantly enriched within the sequences of the 66 TE families with significant regulatory activity (Fig. 7b). In summary, most TE families function as CREs in the primate liver, either as strong enhancers or as repressors. TEs are actively recruited into the regulatory landscape, further supporting our findings on the pervasive involvement of TEs in the primate gene regulation.

## Discussion

### The primate regulatory landscape is evolutionarily conserved

Only a small fraction (<1.50%) of the CREs were differentially active between humans and chimpanzees. This suggests that even modest changes in gene regulation produce observable phenotypic differentiation, and confirm that cis-regulatory evolution plays a central role in primate diversification (Davidson 2001, 2006; Wray, 2007; Ho et al., 2009; Tsankov et al., 2010; Smith et al., 2013; Martin et al., 2012; Coolon et al., 2014; Martin and Reed, 2014; Guo et al., 2015; Lynch et al., 2015; Villar et al., 2015; Adachi et al., 2016; Landeen et al., 2016; Lesch et al., 2016; Zhang and Reed, 2016). Our approach for the comparison of CREs across species, based on the analysis of differential histone modification state in orthologous regions, demonstrated that cis-regulatory divergence across species may be overestimated when based on the peak overlap status as a binary variable.

### Specific genomic features are associated with CRE conservation

Evolutionarily conserved promoters and enhancers have conserved nucleotide sequence, are close to protein coding genes, are functional in many cell types, and harbor many TFBSs. Many regulatory pathways, specifically those involved in the regulation of liver function and housekeeping functions, are strongly conserved across primates, while other pathways, such as immune response, are less constrained and evolve more rapidly. This observation is consistent with the expected arms-race in evolution between host and pathogens.

### Newly evolved CREs are derived from TE exaptation

Based on our findings, exaptation of transposable elements into functional promoters and enhancers is a pervasive phenomenon in primate genomes. LTRs and SVAs are the most frequently exapted in humans and other apes, despite not being among the most common transposable elements in the genome. Primate liver CREs are enriched in young TEs. These young TEs, after being recruited into the primate regulatory network, introduced novel gene expression in the associated species. To our knowledge, this is the first study demonstrating how specific genomic features are associated to the recruitment of TEs as functional elements in the primate regulatory landscape. In contrast, only a minor fraction of the evolutionarily conserved CREs overlap an annotated TE. This suggests the action of strong selection against the disruption of these regulatory elements, in order to maintain stable gene expression. Further, these data suggest that the core regulatory network that establishes liver cell-type identity is very conserved, whereas adaptive evolution occurs on the periphery of the network, where TEs have the most impact on gene regulatory evolution.

## Data availability

All non-human raw sequence data have been deposited in the Sequence Read Archive under following BioProject IDs: PRJNA349047 for RNA-seq and PRJNA349046 for ChIP-seq data.

## Supplemental information

Supplemental can be found with this article online at **xxxxxxxx**

## Author contributions

MT, MC and CDB conceived the project. MT, YP, GHP and CDB designed the taxon sampling and experiments. MT performed ChIP-seq and RNA-seq experiments. MT and MHB performed the parallelized reporter assays. MHB and KA produced luciferase assay data on *GRIN3A* and *JARID2.* KM and VJL designed and performed the luciferase assays on the 69 TE families. MT, VJL and KM performed the TE enrichment analysis. YP designed computational pipelines for the detection of orthologous regions in the Ensembl MSA alignment and all related analyses. MT, YP and CDB analyzed the data and wrote the paper. All authors read and approved the manuscript.

## Acknowledgements

We thank Texas Biomedical Research Institute, Duke University Lemur Center and the Hospital of University of Pennsylvania for providing the samples. We thank Jonathan Schug and other members of UPenn NGS Core Facility. We are grateful to Sarah Tishkoff, Yoseph Barash, Benjamin Voight, Yana Kamberov, Erin Fry, and Barbara Engelhardt for providing insightful comments on the preliminary versions of the results and of the manuscript.

## Materials and Methods

### Tissue sampling

We obtained liver tissue samples for three to four individuals belonging to each of the studied primate species (three bushbabies, four chimpanzees, three humans, three marmosets, three mouse lemurs, and three rhesus macaques; Table S1). Samples for chimpanzees, marmoset and rhesus macaque were obtained from Texas Biomedical Research Institute (San Antonio, TX); bushbaby and mouse lemur livers were obtained from the Lemur Center of Duke University (Durham, NC). Tissue samples were collected and flash-frozen immediately. With the exception of the bushbaby, samples for all of the species included both males and females (Table S1). Age was comparable across species (young adults) and all individuals died of causes unrelated to liver disease.

### RNA-seq sample processing

We processed samples from all species in random batches of four in order to minimize batch effects. For each sample, 25 mg of frozen liver tissue was used to extract total RNA and genomic DNA, using QIAGEN AllPrep DNA/RNA/miRNA Universal Kit. Quality of total RNA was assessed computing the RNA Integrity Number (RIN) using Agilent Bioanalyzer. All RNA samples had a RIN > 8. We used 4µg aliquots of total RNA to produce barcoded RNA sequencing libraries using the Illumina TruSeq Stranded mRNA kit. The quality of generated libraries was assessed using Agilent Bioanalyzer High Sensitivity DNA Kit and Kapa metrics. Libraries were pooled in two different pools based on barcode compatibility, and each pool was sequenced in two Illumina HiSeq2500 lanes, producing on an average of 42.1 million single end (SE) 100-bp reads per sample.

### ChIP-seq sample processing

We processed samples in six randomly assigned groups in order to minimize batch effects. For each sample, we cut 90 mg of frozen liver tissue into 1 mm^3^ pieces, washed the cut tissue samples with cold phosphate-buffered saline (PBS), and fixed with 1% formaldehyde for 5 minutes at room temperature. We prepared nuclei of each washed sample using the Covaris truChIP Tissue Chromatin Shearing Kit. Chromatin was then sheared for 16 minutes using a Covaris S220 Focused-ultrasonicator. We quantified shearing efficiency and chromatin concentration using Agilent Bioanalyzer High Sensitivity DNA Kit.

From each specimen, we kept aside a 0.5 µg aliquot of sheared chromatin to be used as input. We used two 5 µg aliquots of chromatin per sample to perform immunoprecipitation (IP) with antibodies directed at H3K27ac (ab4729) and H3K4me1 (ab8895) respectively. We performed each IP using 5 µg of antibody with an overnight incubation at 4°C as specified by the Magna ChIP A/G Chromatin Immunoprecipitation Kit protocol. After elution and protein-DNA crosslink reversal, we extracted DNA using Zymo Research ChIP DNA Clean & Concentrator kit, and quantified extracted DNA using Agilent High Sensitivity kit and Qubit 2.0.

We used 5 to 15 ng of input and immunoprecipitated DNA to generate sequencing libraries using the NEBNext Ultra ChIPseq library kit, following protocols specified by the manufacturer. We assessed the quality of each constructed library using Agilent Bioanalyzer High Sensitivity DNA Kit and Kapa metrics. Libraries were multiplexed, pooled and sequenced on a total of 16 Illumina HiSeq2500 lanes, producing on an average of 40.6 million SE 100-bp reads per sample.

### Sequence QC: ChIP-seq and RNA-seq

We performed the standard quality control (QC) measures on both ChIP-seq and RNA-seq fastq files using FastQC v0.11.3 (Andrews, 2010). We then trimmed sequencing adapters and low quality base calls using TrimGalore! v0.4.1 with the following parameters: -stringency 5 -length 50 -q 20 (http://www.bioinformatics.babraham.ac.uk/projects/trim_galore/).

### RNA-seq alignment and gene expression quantification

We aligned all sequences that passed QC to the reference genomes from the Ensembl database (bushbaby: otoGar3; chimp: CHIMP2.1.4; humans: GRCh38; rhesus macaque: Mmul1; marmoset: C_jacchus3.2.1; mouse lemur: micMur1) using STAR v2.5 (Dobin et al., 2013) in 2-pass mode with the following parameters: --quantMode TranscriptomeSAM --outFilterMultimapNmax 10 --outFilterMismatchNmax 10 --outFilterMismatchNoverLmax 0.3 --alignIntronMin 21 --alignIntronMax 0 --alignMatesGapMax 0 --alignSJoverhangMin 5 --runThreadN 12 --twopassMode Basic --twopass1readsN 60000000 --sjdbOverhang 100. We filtered bam files based on alignment quality (q = 10) and sorted using Samtools v0.1.19 sort function (Li, 2009). We used the latest annotations for each species obtained from Ensembl to build reference indexes for the STAR alignment: Homo_sapiens.GRCh38.82.chr.gtf; Pan_troglodytes.CHIMP2.1.4.82.chr.gtf; Macaca_mulatta.MMUL_1.82.chr.gtf; Callithrix_jacchus.C_jacchus3.2.1.82.chr.gtf; Otolemur_garnettii.OtoGar3.82.gtf; Microcebus_murinus.micMur1.82.gtf (Aken et al., 2016). We used FeatureCounts (Liao et al., 2014) to count reads mapping to each protein coding gene/lincRNA/pseudogene, according to Ensembl annotations for the six studied species. Read counts were then normalized based on feature (gene) length.

### Differential gene expression analysis

We analyzed differential gene expression levels for each species using read counts, normalized by feature length with the DESeq2 software (Love et al., 2014), and with the following model:

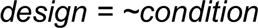

where condition indicates the species or the group of species (e.g. apes).

We used a set of 10,243 genes annotated as orthologs in the six species according to Ensembl (BioMart v. 0.9; Smedley et al., 2015; Table S3) and used 5% False Discovery Rate (FDR) as our multiple-testing-corrected significance threshold. The overall analysis included three main comparisons of our interest: 1) pairwise comparisons between human and each of the other five species; 2) comparisons for human-specific differential expression of orthologous genes (human versus other five primates grouped together); 3) comparisons for ape-specific differential expression of orthologous genes (human + chimpanzee versus other four primates). 4) comparisons for Catarrhini-specific differential expression (human+chimpanzee+rhesus macaque versus other primates). Finally comparison between Strepsirrhini (human, chimpanzee, rhesus macaque and marmoset) and Haplorrhini (mouse lemur and bushbaby) was computed.

### ChIP-seq QC and alignment

We applied standard QC measures to ChIP-seq data as described above for RNA-seq data processing. We aligned the sequences that passed QC to the reference genomes from the Ensembl database (bushbaby: otoGar3; chimpanzee: CHIMP2.1.4; humans: GRCh38; rhesus macaque: Mmul1; marmoset: C_jacchus3.2.1; mouse lemur: micMur1) using Burrows Wheeler Alignment tool (BWA), with the MEM algorithm (Li, 2013). We sorted the filtered bam files using Samtools v0.1.19 (Li, 2009).

### ChIP-seq peak calling and QC

We called peaks for each individual using MACS2 (Zhang et al., 2008), at 1% FDR, and with parameters recommended for histone modifications (Liu, 2014): -- m 30 40 --ext size 147 -B. Input was used as a control in all differential histone modification analyses. We performed QC on peaks called for each specimen using metrics recommended by ENCODE (Landt et al., 2012): Fraction of Reads in Peaks (FRiP), Normalize Strand Correlation coefficient (NSC) and Relative Strand Correlation coefficient (RSC) and ENCODE quality score.

In order to compute FRiP, we used Bedtools (v2.25.0; Quinlan and Hall, 2010) to intersect bed files containing all coordinates of called peaks (narrowPeak output of MACS2) with the original sorted bam file of the specific ChiP-seq sample. Then, we used a publicly available perl script to count the reads mapping in the intersection regions(https://github.com/mel-astar/mel-ngs/blob/master/mel-chipseq/chipseq-metrics/getCnt.pl). As recommended by the ENCODE consortium, we selected a threshold of 1% as acceptable FRiP values. We computed the two strand correlation metrics (NSC, RSC) using Phantompeakqualtools (Landt et al., 2012). For H3K27ac, NSC ≥ 1.05 and RSC ≥ 0.8 were used as threshold for retaining samples. For H3K4me1, that tends to produce broader peaks, we used NSC ≥ 1.05 and RSC ≥ 0.5 (Table S1).

Samples that did not pass at least two of the three main QC metrics (FRiP, NSC, RSC) were excluded for any downstream analysis. We then called human consensus peaks for H3K27ac and H3K4me1 using MACS2 and the above described parameters with the 1% FDR threshold. All human samples passing QC were considered as replicates of each other for the consensus peak calling. These human consensus H3K27ac and H3K4me1 profiles were used to perform all of the below described human-centric downstream analyses. Peaks called for the other species were only used for the above mentioned QC purposes but were not utilized for any of the downstream analyses, with the exception of the chimpanzee consensus peaks (see below). In order to assess how our data compares to known liver related regulatory regions, we overlapped our set of human consensus peaks to the set of H3K27ac and H3K4me1 peaks generated for HepG2 cells from the ENCODE consortium and to the set of permissive enhancers generated by the FANTOM5 consortium.

### Comparison to previous findings using human liver ChIP-seq data

We compared our human H3K27ac ChIP-seq data, with a set of published human liver H3K27ac peaks (Villar et al., 2015). We used the window function of Bedtools to quantify the number of H3K27ac peaks in the replication dataset that either overlap with, or are found within 1 kb up- and downstream from each of our human H3K27ac peaks. With this approach, we quantified and assessed overlaps between discovery and replication datasets.

Next, we characterized sets of replicated and unreplicated peaks between the two datasets. Using the procedures previously described for the comparison of conserved and recently evolved CREs, we annotated several genomic features for both replicated and unreplicated peaks. Specifically, we included information regarding: 1) the average distance from the closest TSS; 2) the class of the associated gene (e.g., protein coding, lincRNA); 3) the number of cell types with an overlapping ENCODE DHS site and; 4) the number of ENCODE TFBS overlapping the peak. Moreover, a logistic regression on the q-values of replicated and not replicated peaks was performed.

### Parallelized reporter assay

We obtained a list of 334 putative 1-kb long CREs overlapping liver eQTLs from Brown and collaborators (Brown et al., 2013). This data included both enhancers (distance from TSS > 1Kb) and promoters (distance from TSS < 1kb). 122 CREs out of these 334 CREs overlapped our human ChIP-seq peaks (53 enhancers and 69 promoters; Table S6). Within each of the loci defined by the investigated liver eQTLs, we predicted a 1-kb CRE. These predicted CREs were amplified in individual PCRs performed on 120 pooled Yoruban HapMap DNA samples. PCR products from each reaction therefore represent a complex mixture of haplotypes. We inserted barcodes (hereafter, tags) consisting of a 160-bp oligo, including a randomized 20-bp unique barcode for each construct, into luciferase reporter vectors (pGL4.23 and pGL4.10), immediately downstream of the luciferase gene, after linearizing the vector with the XbaI restriction enzyme.

We pooled and cloned DNAs from each putative CRE into uniquely barcoded luciferase reporter vectors (pGL4.23 were used for enhancers and pGL4.10 for promoters), using the Gibson Assembly Kit (New England BioLabs). The CREs were specifically inserted upstream of the luciferase gene, after linearizing the vector with the restriction enzymes KpnI and XhoI. We then transfected the complex pool of CRE reporters into HepG2 cells in two replicates. 24 hours after transfection, we extracted total RNA, purified poly-A RNA, and produced cDNA that was used to amplify the tag, with the QIAGEN One Step RT-PCR Kit with primers that included Illumina adapters for sequencing. Tag libraries were pooled and sequenced on a single Illumina HiSeq2500 lane, producing single end (SE) 50-bp reads. We amplified the tags from the vector before the transfection and sequenced them in the same pool with the tag-RNA libraries as a control for tag read counts.

In parallel, reporter tags were unambiguously associated with each specific CRE by sequence based sub-assembly. Briefly, we cut the luciferase gene from the vector by inverse PCR and then re-ligated the vector using the T4 Polynucleotide Kinase (PNK) + T4 ligase kit from NEB. In this way CREs and tags were flanking each other and CRE-tag complexes. The CRE-tag complexes were then PCR amplified using a reverse primer that included Illumina adapter for sequencing. Next, the CRE-tag PCR product was digested for 5 minutes at 55°C using Nexetera Tn5 Transposase (TDE1) in order to produce fragments of variable length (from ca. 150 bp to the entire length of the construct). When cutting the fragments, TDE1 also inserts an Illumina compatible adapter in proximity of the cutting site. We performed a PCR to enrich the libraries using the TDE1 inserted adapter as forward primer and the previously included Illumina adapter as reverse primer.

We pooled the two libraries (one for pGL4.10 and one for pGL4.23 constructs) and sequenced them on an Illumina MiSeq, producing paired-end (PE) reads (250 + 50 bp). After performing QC with FastQC v0.11.3, we aligned the sub-assembly sequences to the human genome (GRCh37/hg19) using BWA mem and the bam files were sorted and indexed with Samtools v0.1.19. Finally, we produced a matrix listing all of the CRE-tag associations. Tags associated with more than one CRE were discarded and not used for further analyses. After attributing each tag to its uniquely associated CRE, we used sequence based tag counts (HiSeq reads), normalized by sequencing depth, to quantify the gene expression level driven by each CRE, and therefore its functionality as enhancer/promoter.

For each CRE, we used a count-based generalized linear model to quantify differential expression between RNA (after transfection) and DNA (before transfection), assuming a Poisson error function:

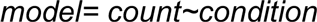

where condition indicates that the read count comes either from RNA (replicates 1 and 2) or DNA-control.

In presence of a significant p-value, the model indicates a significant difference between the expression of the tags in the RNA samples compared to their DNA control. The effect size estimate was then used to infer whether the RNA samples were upregulated, hence showing significantly higher level of expression of the tags compared to their DNA controls, and therefore indicating that the CRE is a functional regulatory element.

### Detection of orthologous regions for human peaks in each primate

We mapped orthologous sequences using all identified consensus ChIP-seq peak regions in both H3K27ac and H3K4me1 experiments. We used the Ensembl multiple sequence alignment (MSA) reference database with the following specifications: 39 Eutherian mammals; method_link_type: “EPO_LOW_COVERAGE”; species_set_name: “mammals” (Herrero et al., 2015). For orthologous sequence analysis, 500 bp up- and downstream regions were considered to be a part of the identified consensus peaks in all six species. We queried all regions directly from the reference database using the REST API (Yates et al., 2015).

All orthologous sequences retained gaps generated by MSA. In cases of incomplete chromosome assembly (e.g. mouse lemur), composite sequence representations containing parts of multiple scaffolds are used as a reference as provided by the Ensembl database. As a result, we independently queried each peak region as well as regions covering each peak +/− 500 bp. All orthologous sequences pulled from the references for downstream analysis contained only directly aligned sequences. All regions with no orthologous regions represented in the MSA reference were excluded from further analyses. All query results in. json format and extracted sequences formatted for the MSA alignment as well as genomic position information are provided in the repository mentioned in the final section.

### Correlation between human and marmoset read counts within orthologous regions

We assessed human and marmoset (i.e. the species with the smallest number of peaks called; Supplemental File S1) normalized read counts at the 47,673 orthologous CREs, after splitting them into two groups: 1) regions with overlapping peaks present in both marmoset and human, and 2) regions with a peak present only in human. Spearman's correlation (ρ) between human and marmoset normalized read depths was then computed for each the two groups.

### Differential histone modification analysis

Using the above described procedure, for both H3K27ac and H3K4me1, we produced a single matrix including the human peaks having an ortholog in each of the studied species and the associated read count for the specific histone mark and for the input in all of the six species. The normalized read counts were used for differential ChIP-seq analysis with DESeq2, performing an interaction analysis using the Wald statistic between the histone marks read counts and their associated input values:

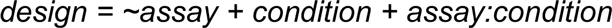

where the assay indicates either histone marks data or input data, and condition indicates instead the species or the group of species (e.g. apes, Catarrhini, Haplorrhini).

Differential histone mark analysis included the same species × species and group × group comparisons described for RNA-seq. We used 10% FDR as our multiple testing corrected significance threshold. Further, different FDRs (5%, 10%, 20%, 30%, 40%) were tested to assess the robustness of our approach.

We analyzed differential histone modifications for the two marks independently. However, in order to quantify the fraction of differentially marked cis-regulatory elements (CREs) in all of the above described pairwise comparisons, we used bedtools to identify the CREs predicted by H3K27ac (i.e. H3K27ac peaks) that would overlap those predicted by H3K4me1 for at least a 25% of their length. The list of unique CREs was used to estimate the fraction of differentially bound CREs for each of the above mentioned human-centric pairwise comparisons. We defined CREs as evolutionarily conserved if they did not show significant differential histone mark in any of the above mentioned pairwise comparisons. Otherwise, the CRE were defined as recently evolved.

### Sequence conservation versus functional conservation analysis

We estimated per-nucleotide pairwise divergence for all five species in comparison to humans using MSA aligned sequences of orthologous regions for consensus peaks +/− 500 bps. All gaps in human were excluded from analysis. Regions not included in the set of six-way orthologous CREs were also pruned. Finally, we removed outliers - with respect to the distribution of the genetic distances in the given pairwise comparison - using the R package *outliers* (Komsta, 2006). We intersected the set of 47,673 orthologous liver CREs with the UCSC phastConsElements30wayPlacental track (Siepel et al., 2005), to assess whether genomic regions characterized by conserved nucleotide sequence (i.e. phastCons elements) are significantly more associated to CREs detected as evolutionarily conserved in primates.

### Analysis of features associated with evolutionary conservation of CREs

The following features were associated to each of predicted human CREs: nearest gene, distance from TSS, functional categories of genes, CRE category and histone mark (Table S2). Any human CREs without orthologous regions in other five species have been excluded from our analyses. To assess the correlation between the degree of conservation of a CRE and the number of cell types where the CRE is functional, we obtained publicly available data for DNase hypersensitivity sites (DHS) for over 200 cell types (ENCODE Project Consortium, 2012). For each CRE overlapping one or more DHS regions, we annotated the number of cell types where the specific CRE is putatively functional.

We estimated the correlation between the degree of conservation of CREs and the number of TFBSs by comparing our human consensus peaks with previously published HepG2 TF-binding profiles (ENCODE Project Consortium, 2012). Further, we quantified the proportion of putative primate CREs overlapping known transcribed enhancers (eRNAs) by using 43,011 known permissive transcribed enhancers (FANTOM5 Consortium, 2014). Similarly, for liver-specificity of human CREs analysis, we used coordinates of published liver eQTLs (Innocenti et al., 2011). We then selected all genes within 10 kb distance from evolutionarily conserved CREs for gene set enrichment analysis using GOrilla software (Eden et al., 2007; Eden et al., 2009). All genes found within 10 kb of any of the 47,673 orthologous CREs are used as a background for the enrichment test.

### Known and *de novo* motif analysis

Genomic coordinates of orthologous regions were used to extract target sequences from the Ensembl references without MSA alignment gaps. All regions containing consensus peaks identified as human- and apes-specific and primate-conserved were used for the motif discovery and enrichment analysis. MEME-chip was used for known motif discovery and enrichment analysis using the Jaspar database (Bailey et al., 2009). We used DREME (Bailey, 2011). De novo motif identification was performed with AME (McLeay et al., 2010). Jaspar and Hocomoco (v10) databases were used as references to estimate similarities to known motifs. All motif discovery and enrichment analysis used default settings and parameters provided by the developers except for the maximum *de novo* motif discovery threshold (changed from 1 to 1000 for maximum threshold). Shuffled input sequences were used to estimate the background distribution of motifs.

### Overlap of transposable elements (TEs) with primate CREs

We used Bedtools to overlap the RepeatMasker track for GRCh38 to the set of unique human CREs that would overlap a TE for at least 25% of their length. TE enrichment analysis was performed using the *TEAnalysis* pipeline with TE-analysis_Shuffle_bed v. 2.0, setting 1000 replicates (https://github.com/4ureliek/TEanalysis; Kapusta et al., 2013). To test the regulatory effect of enriched young TEs on primate gene expression, we performed a differential gene expression analysis between Strepsirrhini (human, chimp, rhesus macaque, marmoset) and Haplorrhini (bushbaby, mouse lemur) and quantified the number of CREs associated to differential expressed genes that overlapped a TE younger than the Haplorrhini-Strepsirrhini divergence.

### Analysis of SINE-VNTR-*Alus* (SVA) TEs enriched in human CREs

We intersected all known SVAs annotated in the human genome with all human consensus peaks from our study. We used the same 25% overlap threshold as described above and considered all human consensus peaks regardless of presence of orthologous regions in the other species. The two SVA lists (overlapping and not overlapping the CREs, respectively) were annotated for: the average distance from the closest TSS, functional categories of associated gene, the average number of cell types with available DHS data, the average number of TFBS overlapping the SVAs, and finally the average number of TFBS within 10 kb up- and downstream of the SVAs. We used AME (McLeay et al., 2010) to look for motifs enriched in the exapted SVAs, using the not exapted SVAs as a control.

### Luciferase reporter assay validation of *GRIN3A* and *JARID2*

To test for species- or clade-specific regulatory activity, we compared activity of two predicted functional CREs with the empty pGL4.23 vector as a negative control. For *GRIN3A* we PCR amplified the CRE (Table S6), and cloned the fragment into pGL4.23 using the NEB Gibson Assembly Kit. The *JARID2* CRE was synthesized by GenScript and cloned into the same pGL4.23 vector. Cells were grown in DMEM high glucose (Gibco #11965084) supplemented with 10% fetal bovine serum (FBS) (GE Healthcare Life Sciences #SH3091003) containing antibiotic and antimycotic (Gibco #15240062) in a humidified incubator with 5% CO_2_ at 37°C. HepG2 cells were seeded in 48-well CellBIND surface plates (Costar #3338) with 1.5 x 10^5^ cells per well 24 h prior to transfection. Transfection complexes were formed using 800 ng of each construct with 1 μL of TransIT-LT1 transfection reagent (Mirus #MIR2304) and Opti-MEM (Gibso #31985070) in a total volume of 27 μL, incubated for 20 min and then added to cells. After transfection, cells were incubated for 24 h and were lysed in passive lysis buffer. To read firefly luciferase activity, 100 μL of LARII were added to 20 μL of cell lysate (from the dual-luciferase reporter assay system from Promega #E1910). We read Luminescence for 2 seconds per well on a 96-well compatible plate luminometer (ThermoFisher Luminoskan Ascent). The constructs were tested using three vector preparations in three to four technical transfection replicates (9 to 12 measurements for for each construct). We normalized for transfection replicates effect using a linear model:

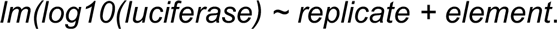

### Validation of the gene regulatory functionality of TE families

HepG2 cells were cultured in DMEM + GlutaMAX (Gibco) supplemented with 10% Fetal Bovine Serum (Gibco) and Normocin (InvivoGen). Transposable element constructs were built by synthesizing (GenScript) the Dfam (Hubley et al., 2016) consensus sequence for each element and cloning into the pGL3 Basic vector (Promega) with an added minimal promoter (pGL3 Basic[minP]). pGL3 BASIC[minP] with no insert was used as the negative expression control. pRL null (Promega) was the renilla control for transfection efficiency. TAP2 cloned into the pGL3 Basic[minP] was the positive control. Confluent HepG2 cells in opaque 96 well plates in 90ml of Opti-MEM (Gibco) were transfected according to the Lipofectamine p3000 protocol (Invitrogen) with 100 ng of the luciferase containing plasmid, 1 ng of pRL null, 0.3 ml of Lipofectamine 3000, and 0.2ml of p3000 reagent in 10 ml of Opti-MEM per well. The cells incubated in the transfection mixture for 24h hours then the media was then changed to the regular FBS containing media for an additional 24 hours. Dual Luciferase Reporter Assays (Promega) were started by incubating the cells for 15 mins in 20 ml of 1x passive lysis buffer. Luciferase and renilla expression were then measured using the Glomax multi+ detection system (Promega). Luciferase expression values of the transposable elements and TAP2 were standardized by the renilla expression values and background expression values as determined by pGL3-Basic expression. Enriched motifs were found by analyzing the Dfam (Hubley et al., 2016) consensus sequences of the TEs found to have a regulatory ability significantly different from the pGL3 Basic[minP] empty vector using the MEME Suite. TomTom (Gupta et al., 2007) was used to match binding site motifs in the Jaspar database to the enriched motifs found in our data.

### Additional notes on analyses used throughout the project

All statistical analyses (DESeq2 analysis, Fisher’s exact tests, logistic regressions, Spearman’s correlations, Wilcoxon tests, General Linear Models, binomial test, and quantiles calculations) were performed using R v3.3.1. Figures were made with the package ggplot2 (Wickham, 2009) in R v3.3.1. Bedtools v2.25.0 (Quinlan et al., 2010) was used for overlap and closest feature/window analyses. All relevant scripts and pipelines are available online (https://github.com/ypar/cre_evo_primates.git). All supplementary data are also available online (https://github.com/ypar/cre_evo_primates_data.git).

---

**Figure S1:**
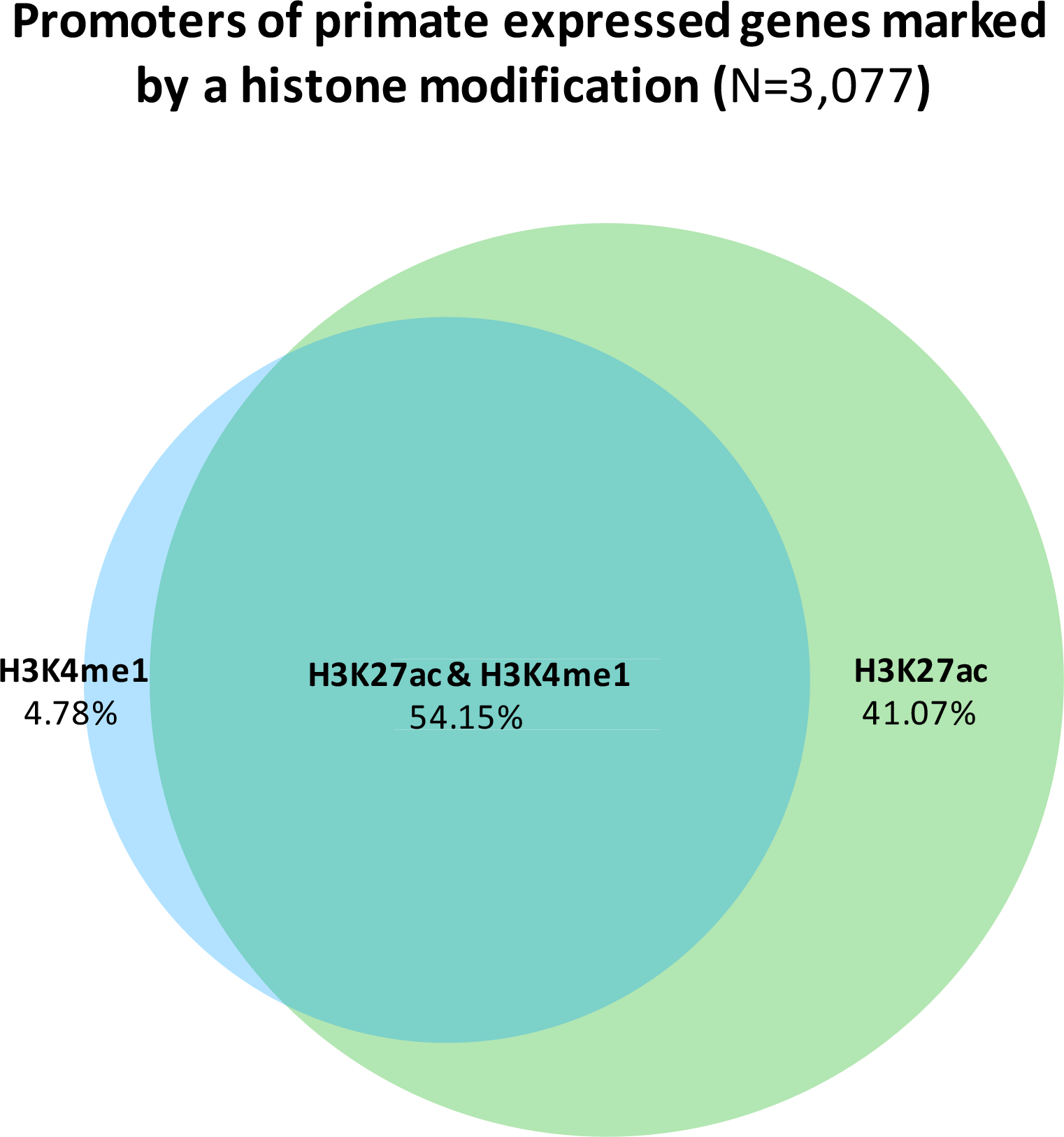
Promoters of genes that are expressed in primates are significantly histone modified. Venn diagram showing the distribution of histone marks on the promoters of genes that are expressed in the primate liver.

**Figure S2:**
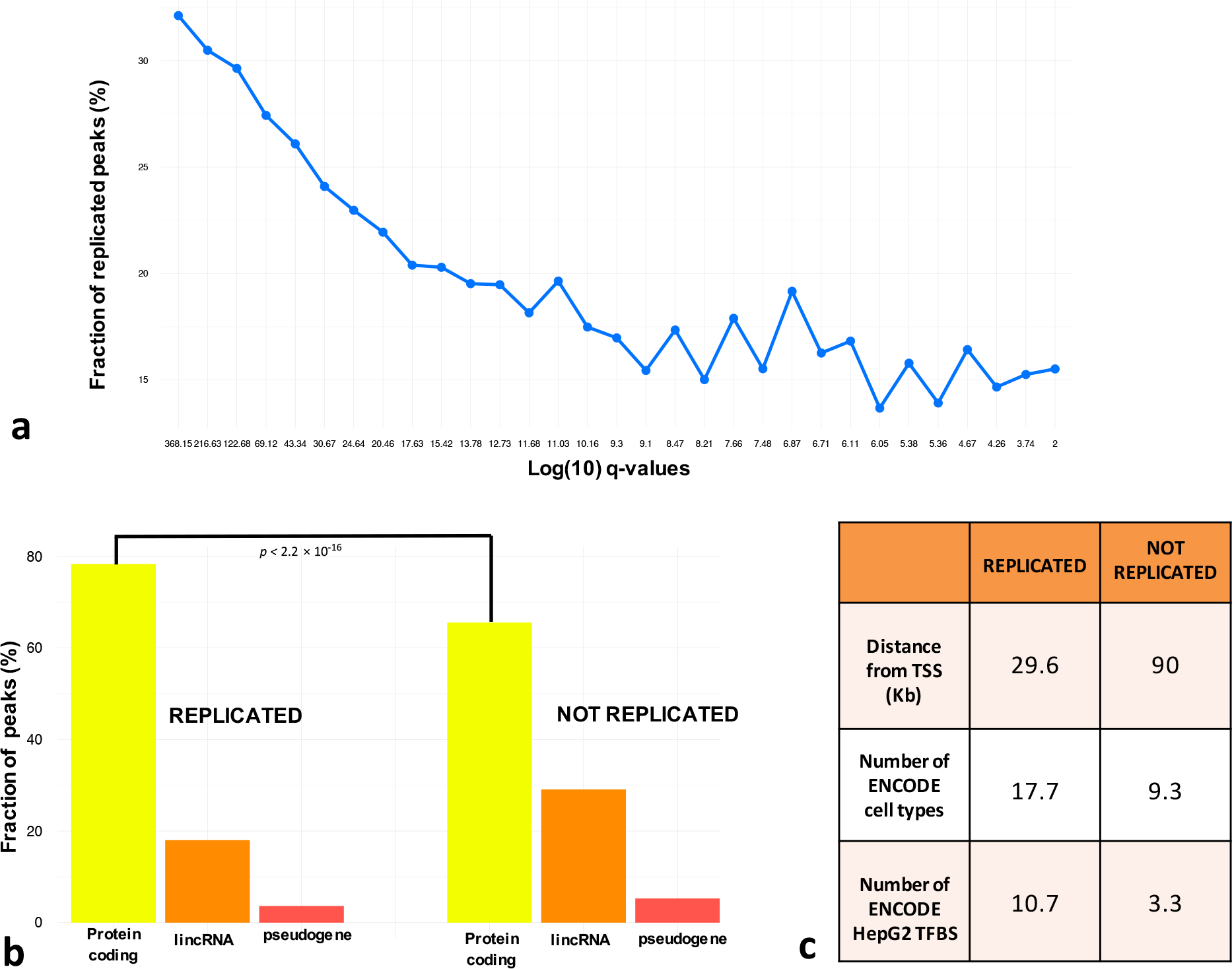
Peaks bearing signatures of robust and broad regulatory activities are largely reproducible across studies. We compared our human H3K27ac data to a recent study focused on liver CREs in mammals (Villar et al., 2015). Overall, slightly less than 40% of the peaks identified by Villar et al. (2015) overlapped one of our H3K27ac regions. Further, 69.3% of the ENCODE HepG2 H3K27ac peaks were replicated in the Villar et al. (2015) dataset. We thus investigated possible features associated with ChlP-seq peak reproducibility. (A) Peak discovery significance (q-value) is significantly correlated with cross-dataset reproducibility (logistic regression *p* < 2.2×10^−16^). (B) Replicated peaks are significantly more likely than non-replicated peaks to be associated with protein coding genes rather than with lincRNAs or pseudogenes (Fisher’s exact test *p* < 2.2×10^−16^). (C) Replicated peaks are: 1) systematically closer to the nearest TSS (29.6 kb for replicated peaks, 90.8 kb for not replicated; Wilcoxon rank-sum test *p* < 2.2×10^−16^); 2) overlap chromatin accessible regions in significantly higher numbers of ENCODE cell types (an average of 17.7 cell types for replicated peaks and 9.3 cell types for unreplicated peaks; logistic regression *p* < 2.2×10^−16^); 3) contain a significantly higher number of transcription factor binding sites (TFBSs) per peak region as identified by ENCODE in HepG2 cells (10.7 TFBSs in the replicated peaks, 3.6 in the unreplicated; logistic regression *p* < 2.2×10^−16^).

**Figure S3:**
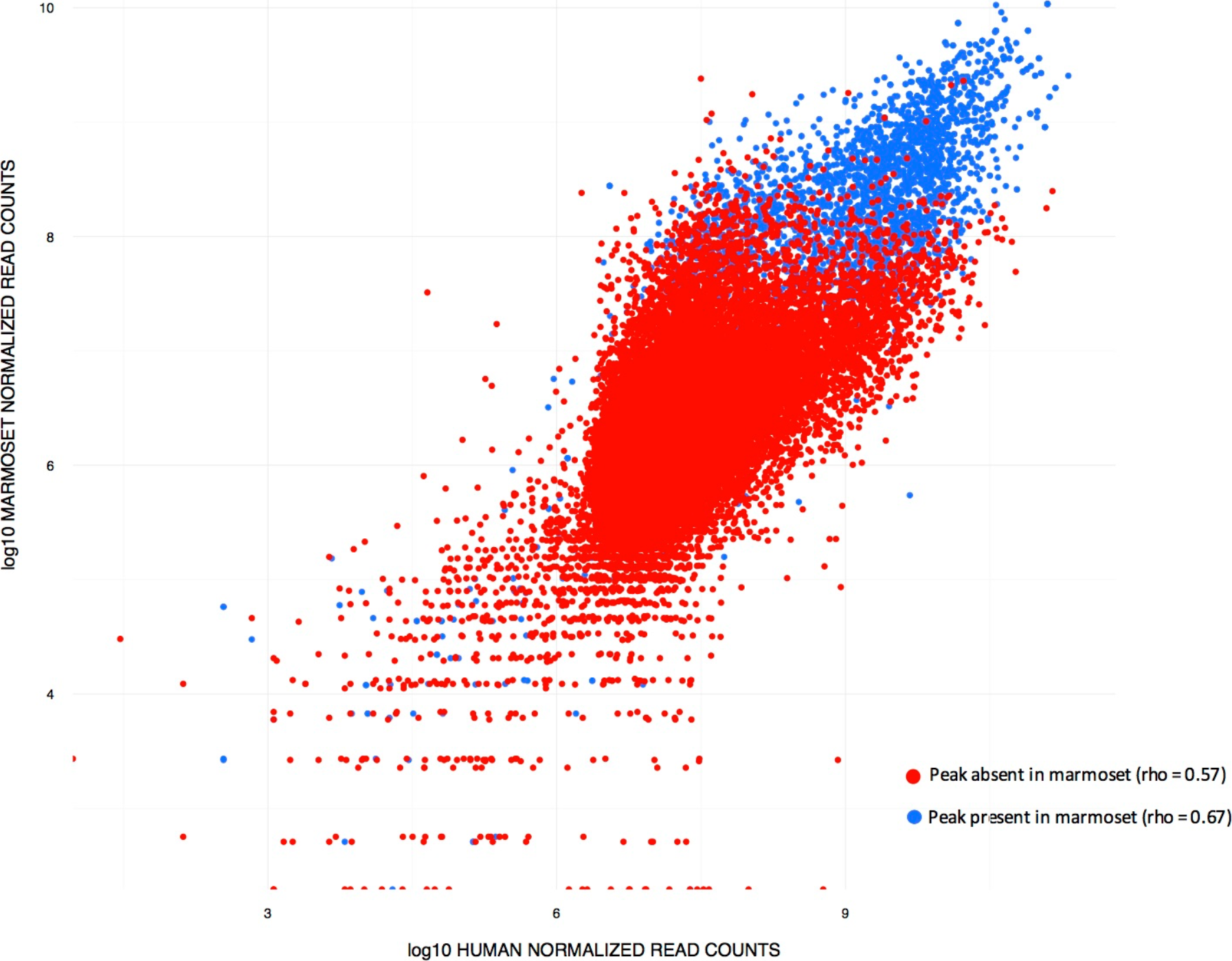
Correlation of the normalized ChIP-seq read depths between human and marmoset. Human consensus peaks with orthologous in all the six species were split into two groups for this analysis: 1) regions with overlapping peaks present in both marmoset and human, and 2) regions with a peak present only in human. While human and marmoset normalized read counts were more highly correlated with each other in group 1 (Spearman’s ρ = 0.67; *p* < 2.2×10^−16^), we found a nearly as strong correlation in group 2 (Spearman’s ρ = 0.57; *p* < 2.2×10^−16^). These findings are consistent with the results of our differential histone modification state analyses, which demonstrated that only a small fraction of the 47,673 orthologous CREs (5.97%, FDR < 10%) are differentially modified, despite the fact that we had a much smaller total number of peak calls in the marmoset samples.

**Figure S4:**
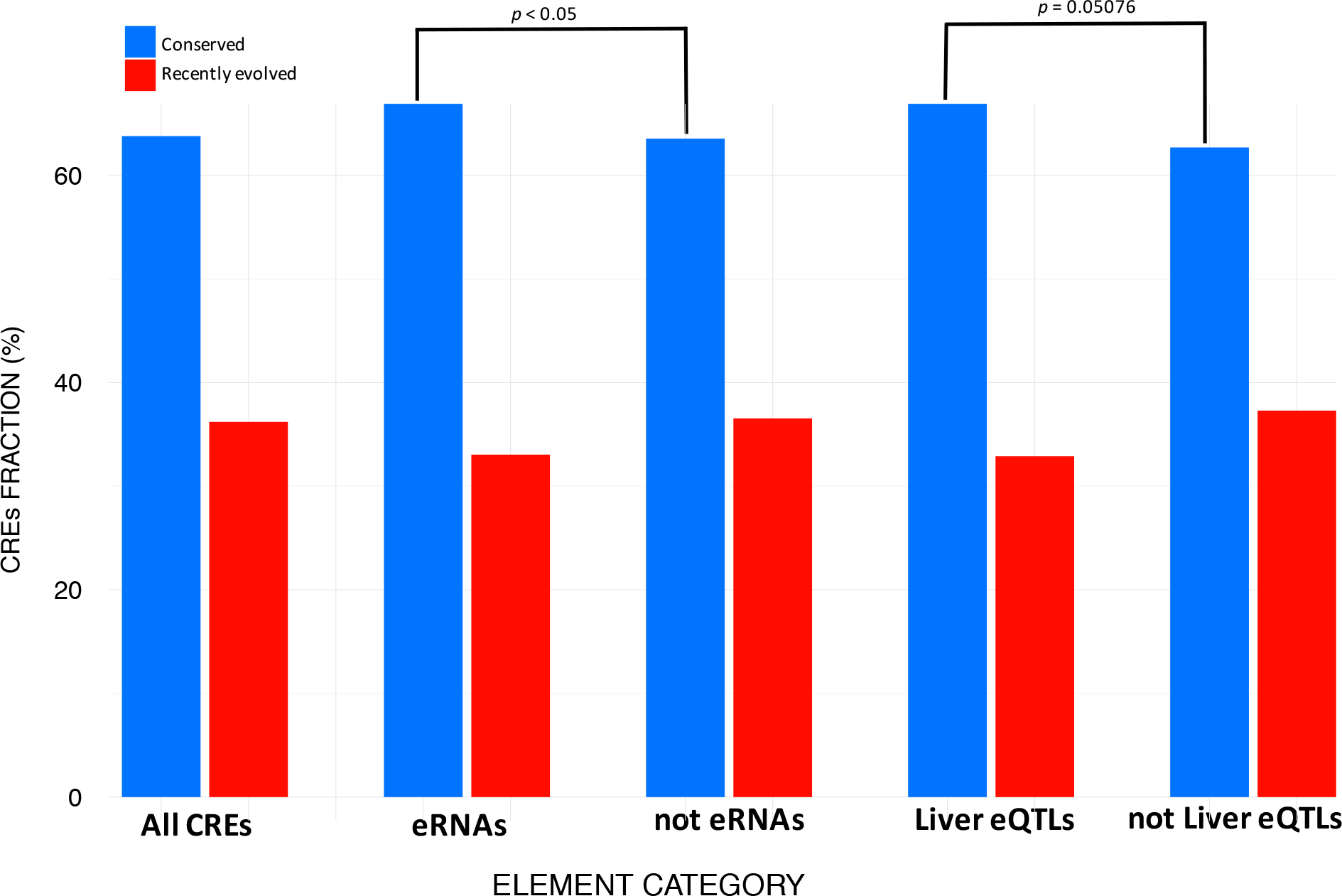
Enhancers with signature of transcription are conserved. Fraction of conserved and recently evolved CREs with signature of transcription (eRNA) based on FANTOM5 Consortium data. Fraction of conserved and recently evolved CREs with no signature of transcription (eRNA) based on FANTOM5 Consortium data, Fraction of conserved and recently evolved CREs overlapping and not overlapping liver eQTLs. Specifically, predicted human CREs overlapped 500 eQTLs detected in a recent study on human liver (Innocenti et al., 2011). We tested whether enhancers and promoters overlapping liver eQTLs would lean toward being more conserved or more labile than a random liver CRE, and we found that neither of these two conditions are satisfied (Fisher’s Exact Test *p* = 0.05076), as we show that liver eQTLs behave as “average” liver regulatory elements.

**Fig S5:**
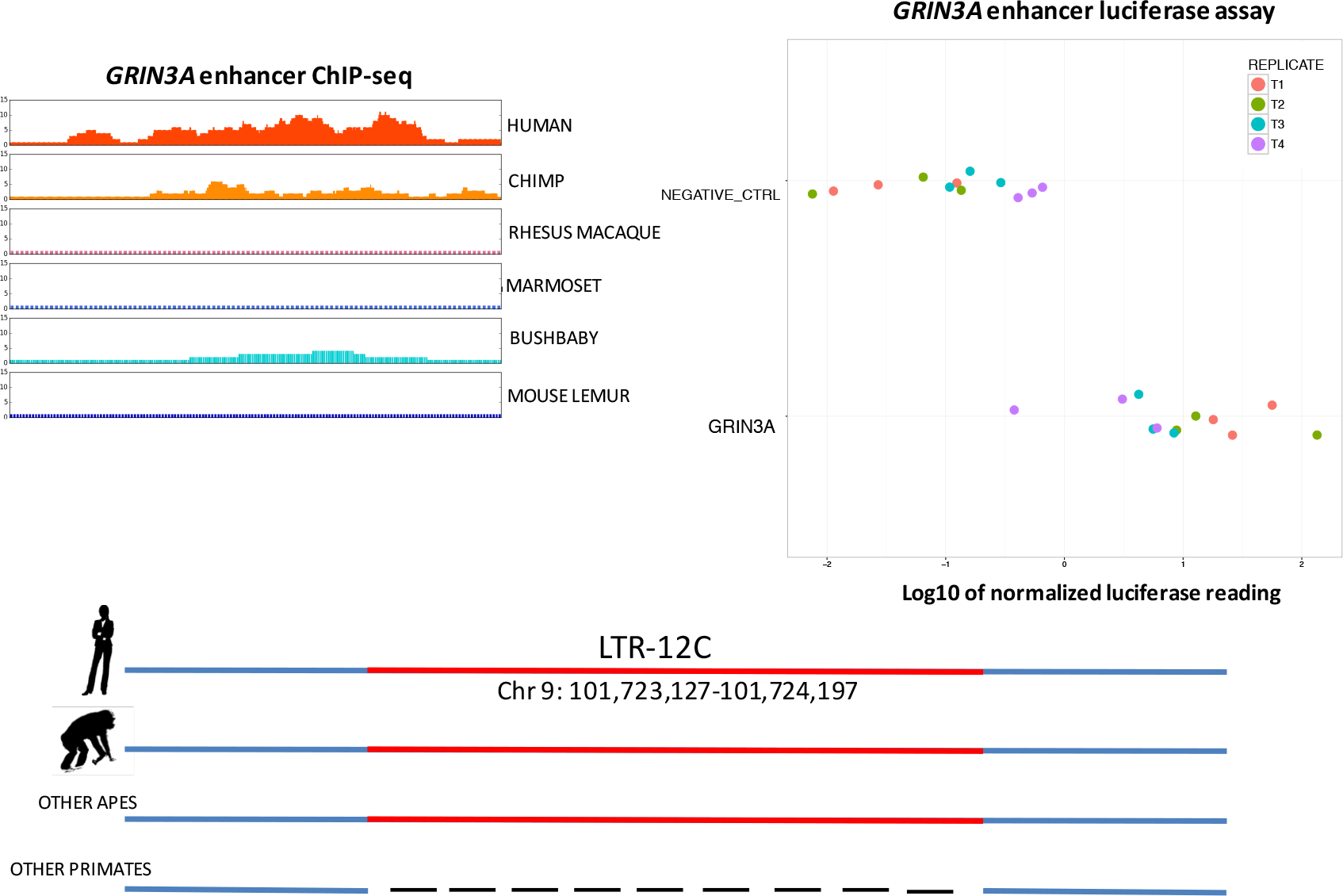
Functional analysis on *GRIN3A locus.* ChlP-seq read depth distributions and luciferase assays reporter activity for the CRE associated to *GRIN3A.*

**Fig S6:**
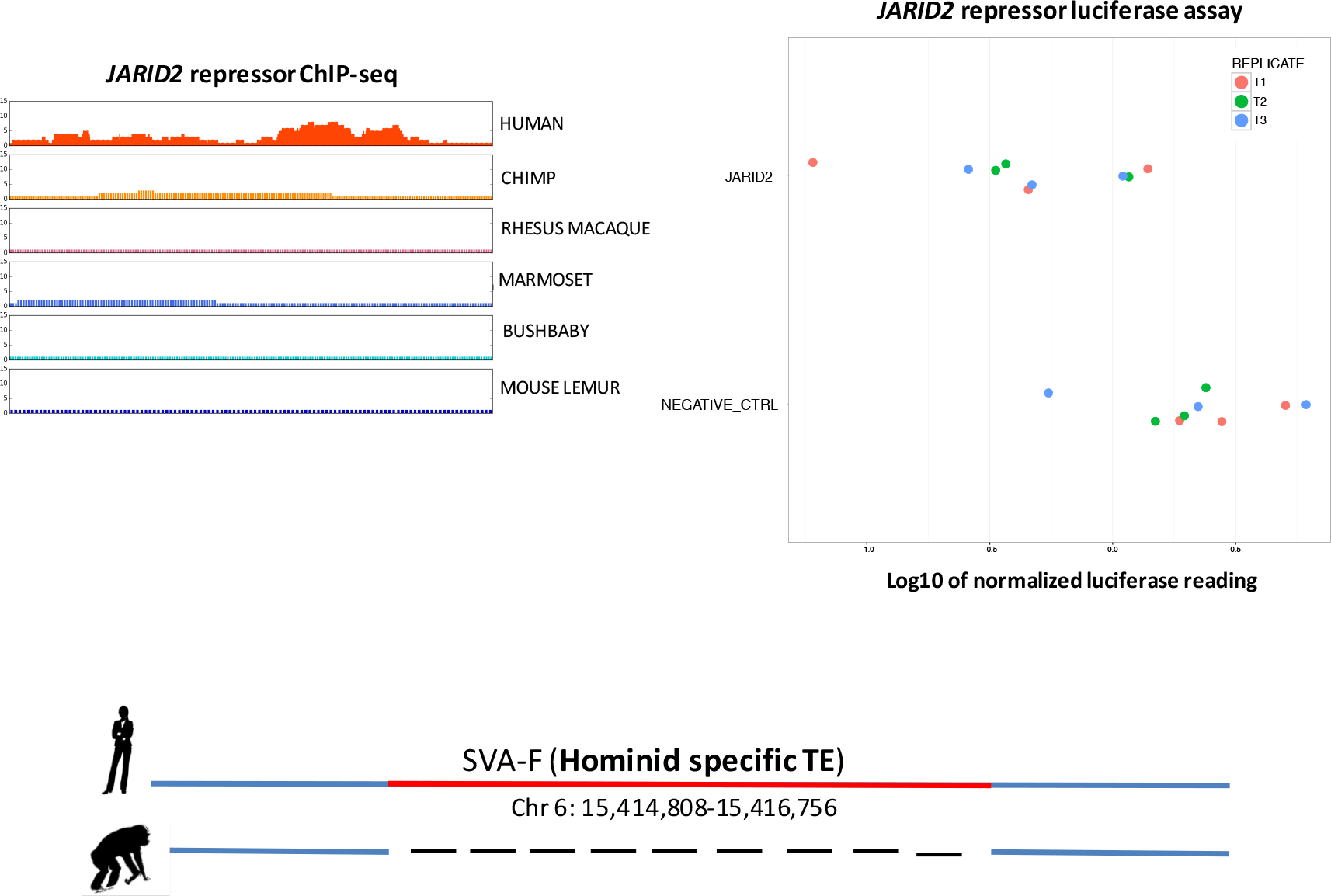
Functional analysis on *JARID2 locus.* ChIP-seq read depth distributions and luciferase assays reporter activity for the CRE associated to *JARID2*.

**Figure S7:**
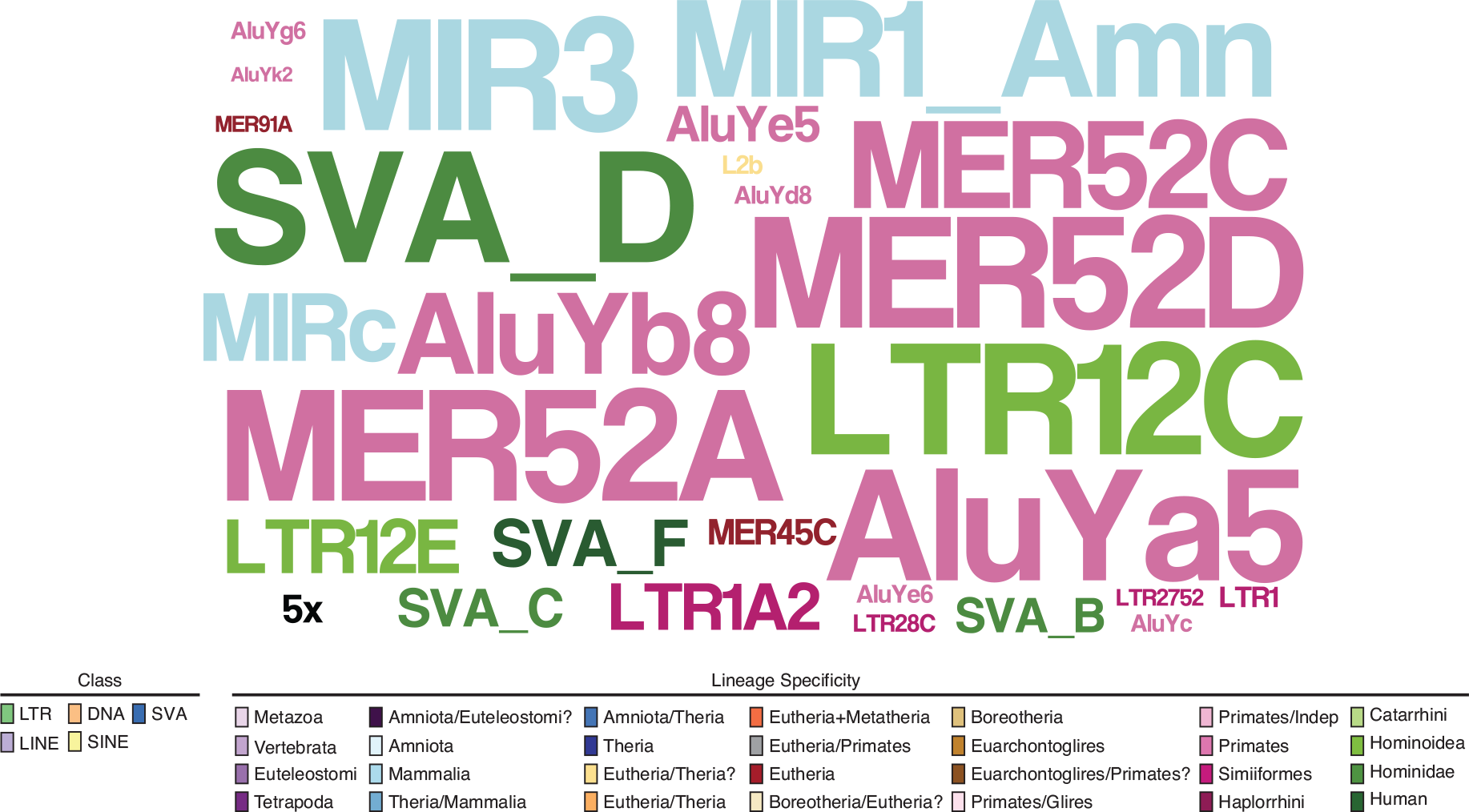
Lineage specificity of enriched TEs. Word-cloud representing the lineage specificity of the enriched TE families.

